# Loss of coiled-coil protein Cep55 impairs abscission processes and results in p53-dependent apoptosis in developing cortex

**DOI:** 10.1101/2020.06.02.129346

**Authors:** Jessica N. Little, Katrina C. McNeely, Nadine Michel, Christopher J. Bott, Kaela S. Lettieri, Madison R. Hecht, Sara A. Martin, Noelle D. Dwyer

## Abstract

To produce a brain of normal size and structure, embryonic neural stem cell (NSCs) must tightly regulate their cell divisions. Cerebral cortex NSCs undergo a polarized form of cytokinesis whose regulation is poorly understood. Cytokinetic abscission severs the daughter cells and is mediated by the midbody at the apical membrane. Here we elucidate the role of the coiled-coil midbody protein Cep55 in NSC abscission and brain development. A knockout of Cep55 in mice causes microcephaly with reduced NSCs and neurons, but relatively normal body size. Fixed and live analyses show NSCs lacking Cep55 have decreased but not eliminated ESCRT recruitment, and have abnormal abscission and higher rates of failure. P53-mediated apoptosis is greatly increased in the brain, but not other tissues, and p53 knockout partly rescues brain size. Thus, loss of Cep55 causes abscission defects and failures in multiple cell types, but the secondary p53 response and apoptosis is brain-specific.

## Introduction

Embryonic neural stem cells (NSCs) must undergo rapid divisions within strict developmental time windows to produce numerous daughter cells of various fates, to build a brain of correct size and structure. They do so within a polarized epithelium, with their apical membranes forming the ventricle wall, and their basal processes contacting the pia. Their nuclei move to the apical membrane for mitosis and cytokinesis. We previously showed cytokinesis in NSCs is polarized and developmentally regulated (McNeely & Dwyer, 2020). The cleavage furrow ingresses asymmetrically, forming the midbody within the apical membrane. The midbody then mediates severing of the intercellular bridge, in a process called abscission. Errors in cytokinesis are one etiology of microcephaly, in which the brain is disproportionately small relative to body size (Bondeson et al., 2017; Di Cunto et al., 2000; Frosk et al., 2017; Li et al., 2016; Makrythanasis et al., 2018; Moawia et al., 2017). We previously showed mutation of the Kinesin-6 family member Kif20b disrupts normal cytokinetic abscission to cause microcephaly in mice (Dwyer et al., 2011; Janisch et al., 2013; Little & Dwyer, 2019). The unique constraints of NSCs during cell division have been hypothesized to make brain growth more vulnerable to defects in mitosis or cytokinesis than other tissues. However, the cellular and developmental mechanisms that drive these tissue-specific requirements are just beginning to be identified.

Cytokinesis has long been studied in cell lines and invertebrate organisms; this work has identified many required proteins, and generated a model for the mechanism of abscission (Green et al., 2012; Mierzwa & Gerlich, 2014). Beginning in anaphase following chromosome segregation, the ingressing furrow compacts the central spindle microtubules and associated proteins into a structure called the midbody, within the intercellular bridge. The midbody has a thick central bulge flanked on either side by dense microtubule bundles. This complex structure consists of over 450 proteins that mediate midbody formation, maturation, and the final scission event, in which the midbody flanks are severed to separate the two daughter cells (Addi et al., 2020; Hu et al., 2012; Skop et al., 2004). The abscission process involves both microtubule disassembly and membrane severing, thought to be mediated by the endosomal sorting complexes required for transport (ESCRT) filaments (Connell et al., 2009; Guizetti et al., 2011). The duration of abscission can be regulated, and slower abscission has been linked to stemness (Chaigne et al., 2019; Lenhart & DiNardo, 2015; McNeely & Dwyer, 2020). After abscission, the central bulge domain remains intact as the midbody remnant (MBR). Studies in mammalian cell lines suggest that the MBR may mediate cell signaling and fate, either by binding cell surface receptors or through uptake into daughter or neighbor cells (Crowell et al., 2014; Peterman et al., 2019). We previously showed that in embryonic brains, MBRs can be visualized on the apical membranes of cortical NSCs. Interestingly, MBRs are more persistent at early proliferative stages, and downregulated later (McNeely & Dwyer, 2020).

The coiled-coil scaffolding protein Cep55 is thought to be essential for abscission in cell lines (Fabbro et al., 2005; Zhao et al., 2006), with the key role of initiating the cascade of ESCRT recruitment in midbodies (Stoten & Carlton, 2018). However, invertebrates lack a *Cep55* orthologue in their genomes, yet complete abscission. Human mutations of *Cep55* cause a variety of brain malformations (Barrie et al., 2020; Bondeson et al., 2017; Frosk et al., 2017; Rawlins et al., 2019). That human embryos with Cep55 mutation are able to develop at all is surprising, given that knockdown of Cep55 in human cell lines caused almost universal failure of cell division (Fabbro et al., 2005; Zhao et al., 2006).

Here, we use a mouse knockout to elucidate the roles and requirements of Cep55 in abscission of neural stem cells during brain development. We find *Cep55* is not absolutely required for abscission in NSCs or other embryonic cells, but its loss causes abnormalities and increases failures. NSCs are especially vulnerable to loss of Cep55 compared to other tissues, and brain size is disproportionally reduced. Surprisingly, we find ESCRT recruitment is not eliminated in *Cep55 −/−* cell midbodies, though it is impaired. We use fixed and live imaging of intact cortical epithelium to probe Cep55’s role in abscission. Additionally, we find the severe effect on the nervous system compared to other tissues appears to be due to a specific p53 response triggering apoptosis, and test this using double knockout of Cep55 and p53. This work underlines that studying cytokinetic proteins and processes in the context of developing tissues is necessary to complement single cell models, in order to understand both the cellular roles of specific proteins, and how their loss can cause various phenotypes at the tissue and organism level.

## Results

### *Cep55* knockout results in microcephaly with thin neuronal and axonal layers

The mutation of *Cep55* used in this analysis is a 600 base-pair deletion encompassing all of exon 6 and flanking intronic sequence, generated by the CMMR (see Methods). The total murine *Cep55* gene consists of 9 exons encoding a protein of 462 amino acids (AA) **(Figure 1A).** Cep55 protein domains include two coiled-coil regions (CC1 and CC2) surrounding the ESCRT- and Alix-binding region (EABR), and two ubiquitin binding domains (UBD) in the C-terminus (Lee et al., 2008; Morita et al., 2007; Said Halidi et al., 2019). This deletion is predicted to result in a frameshift starting at AA 227, resulting in multiple premature stop codons starting after AA 237. The Cep55 protein is undetectable by immunoblot of homozygous mutant tissue lysates **(Figure 1B).**

**Figure 1:**
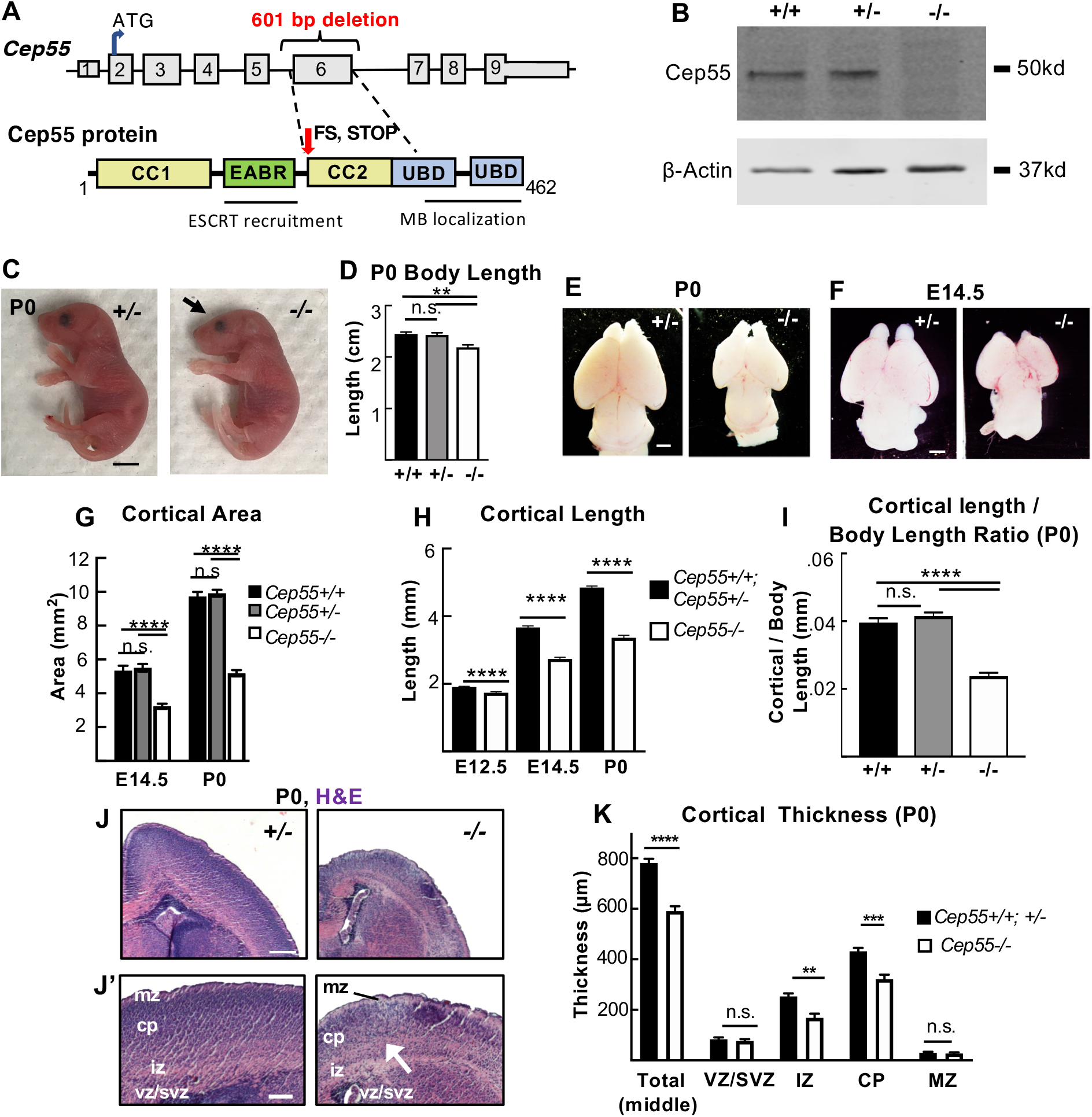
*Cep55* knockout causes microcephaly with severe thinning of neuronal and axon layers. **(A)** Schematics of *Cep55* mouse gene and protein (not to scale). The 601 base-pair deletion includes all of exon 6 (311 bp) and portions of flanking introns. Cep55 protein domains include two coiled-coil regions (CC1 and CC2) surrounding the ESCRT- and Alix-binding region (EABR), and two ubiquitin binding domains (UBD). The deletion of exon 6 is predicted to result in a frameshift (FS) at amino acid 226 which causes multiple premature stop codons starting 12 residues later. (**B)** Immunoblots of E14.5 mouse embryonic fibroblast (MEF) lysates show the expected Cep55 protein product at approximately 55 kDa in wild-type and heterozygote samples, but not in ko samples. **(C,D)** Newborn *Cep55−/−* pups have 10% shorter body length and a flatter head (arrow) but otherwise appear morphologically similar to controls. **(E-G)** Representative images and measurements show mean cortical area is reduced in *Cep55−/−* brains at P0 and E14.5 compared to *+/+* and *+/−* controls. **(H)** Mean cortical length is significantly reduced in *Cep55−/−* at E12.5, E14.5, and P0. **(I)** The cortical length in *Cep55−/−* P0 pups is disproportionately small relative to body size. (**J)** Representative images of P0 *Cep55+/−* and *−/−* coronal sections at level of corpus callosum, stained with H&E. mz, marginal zone; cp, cortical plate; iz, intermediate zone; vz/svz, ventricular/subventricular zone. *Cep55−/−* have reduced cell density in neuronal layers (J’, arrow). **(K)** Mean total cortical thickness is significantly reduced in *Cep55 −/−* brains at P0. The VZ/SVZ and MZ thickness are unaltered, but the intermediate zone (IZ) and cortical plate (CP) are significantly thinner. For P0, n = 7 *Cep55+/+,* 14 *+/−* and 8 *−/−* mice. For E14.5, n = 5 *Cep55+/+*, 13 *+/−* and 10 *−/−* mice; for E12.5, n = 17 Cep55*+/+*;*+/−* and 9 *−/−* mice. For (L), n = 3 *Cep55+/+*, 5 *+/−*, and 5 *−/−* mice. Scale bars: C: 5 mm; E and F: 1 mm; J, 250 μm, J’ 100 μm. n.s., not significant. ** p < 0.01; *** p < 0.001, **** p < 0.0001. D,G, and I: One-way ANOVA; H, K: Student's t-test.

Given that Cep55 was reported to be essential for mammalian cell division (Fabbro et al. 2005; Zhao et al. 2006), we were surprised to discover that *Cep55 −/−* mice are born alive at expected Mendelian ratios, and appear grossly normal with a slightly small body **(Figure 1C)**. However, while the pups survive postnatally, they fail to thrive and die before or around weaning **(Figure 1 – figure supplement 1A-C)**. Upon inspection, *Cep55 −/−* pups have slightly flattened heads **(Figure 1C, arrow; Figure 1 – figure supplement 1D),** and a 10% reduction in body length **(Figure 1D**). Removing the brains from the heads revealed that *Cep55* −/− mice have severe microcephaly, with a 50% reduction in cortical area and 30% reduction in cortical length on postnatal day 0 (P0) **(Figure 1 E, G, H)**. Dissection of prenatal brains showed a similar reduction in brain size compared to controls at embryonic day (E)14.5 and a smaller (8%) reduction at age E12.5 **(Figure 1F-H)**. Cortical size is disproportionately reduced compared to body size **(Figure 1I)**. However, eye size is normal in *Cep55 −/−* pups **(Figure 1 – figure supplement 1D-F)**. In addition to reduced brain size, *Cep55 −/−* mice have reduced cortical thickness at P0 **(Figure 1J, K).** Thus, *Cep55* loss in mice appears to more drastically disrupt brain development than body development, and the reduction in cortical growth correlates with the onset of neurogenesis around E12.5.

To determine if *Cep55 −/−* mice have normal brain structure, we analyzed P0 cross sections stained with hematoxylin and eosin (H&E) to label nuclei and axons, respectively. We noted a reduction in size of all brain regions, including the forebrain, midbrain, and hindbrain **(Figure 1 – figure supplement 2A)**. However, the forebrain appeared to be most severely affected. Interestingly, while cortical thickness is reduced globally, the caudal cortex is the most severely affected **(Figure 1 – figure supplement 2B, C)**.

To begin to determine the cellular cause of the thinner cortex in *Cep55* knockouts, we compared the thicknesses of individual cortical layers in P0 control and mutant brains. Cortical stem (NSC) and progenitor cells reside in the ventricular zone (vz) and subventricular zone (svz), while post-mitotic neurons reside in the cortical plate (cp), and project their axons in the intermediate zone (iz). The cp and iz of *Cep55 −/−* cortices are significantly reduced in thickness, while the vz and svz are not **(Figure 1J, K)**, suggesting a deficit of neurons. To examine the neuronal layers within the cortical plate, we labeled control and *Cep55* −/− cortical sections with Ctip2, for layers 5 and 6, and Satb2, for layers 2-4 **(Figure 1 – figure supplement 2D)**. *Cep55* −/− cortical layers are ordered similarly to controls, but all are reduced in thickness (**Figure 1 – figure supplement 2E)**. As a proportion of the total cp, the deep layers 5-6 of *Cep55 −/−* brains occupy more space, while the upper layers 2-4 occupy less **(Figure 1 – figure supplement 2F)**. Interestingly, both the absolute number and the density of nuclei in layer 6 are significantly reduced **(Figure 1 – figure supplement 2G, H)**. Together these data show the reduced thickness of *Cep55* −/− cortices at birth is attributable to reduced thickness of all neuronal layers and reduced number of neurons.

### NSCs are reduced and cortical layers are disorganized in*Cep55* knockout brains

To begin to address the developmental defects leading to smaller brain size at birth, we examined earlier stages of cortical development. Because *Cep55* −/− brains are already dramatically reduced in size at E14.5, and this is an age when the cortical plate is forming, we analyzed numbers and positions of NSCs, basal progenitors (BPs), and neurons at this age. By labeling E14.5 sections with neuronal tubulin (Tubb3), we noted the cp was already thinner in *Cep55 −/−* cortices, and the iz layer of axons was not distinct **(Figure 2A)**. *Cep55 −/−* cortical thickness is reduced by 12% at E14.5 (compared to 25% at P0, **Figure 1K**), and similar to newborn brains, this reduction is due to thinner neuronal (cp and iz) rather than proliferative layers (vz and svz) **(Figure 2B).** As a proportion of total thickness, the proliferative layer is increased **(Figure 2C)**. These data show that neuron production is already compromised before E14.5.

**Figure 2:**
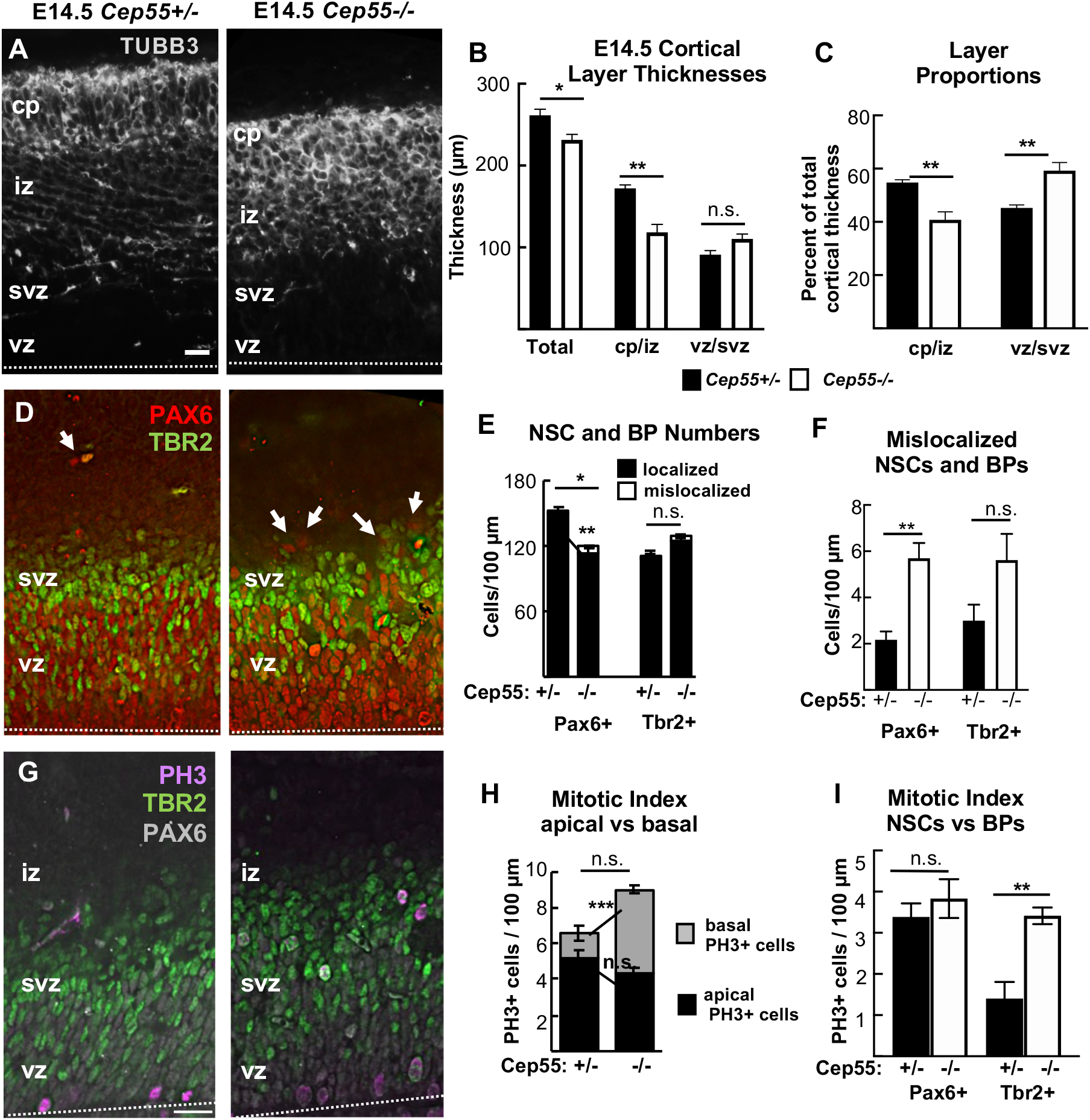
*Cep55* knockout cortex at E14.5 shows reduced and disorganized neuron and NSC layers. **(A)** Representative images of cortical sections from *Cep55 +/−* and *−/−* E14.5 embryos immunostained for neuron marker Tubb3 (gray) show cortical plate (cp) and axons in the intermediate zone (iz). In *−/−* images, the cp is disorganized the border between cp and iz is unclear. **(B)** Mean thicknesses of total cortex and neuron layer are significantly decreased in *−/−* brains. **(C)** The *−/−* cp/iz occupies proportionally less of cortical width than normal, while the vz/svz occupies more**. (D)** Cortical sections stained for NSC marker Pax6 (red) and basal progenitor (BP) marker Tbr2 (green) show NSCs in the vz of *Cep55−/−* brains are more disorganized, with some empty spaces (square), and some nuclei mislocalized basally above the svz (arrows). **(E-F)** NSCs (Pax6+) per cortical length are reduced in *Cep55−/−* and some mislocalized. BP numbers (Tbr2+) are not significantly changed. For B, total thickness, n = 7 *+/−* and 8 *−/−* brains. For B, cp/iz and vz/svz thickness, and for C,E,F, n= 4 *+/−* and 4 *−/−* brains. **(G)** Phospho-histone H3 (PH3) immunostaining is used to mark cells in mitosis. **(H)** *Cep55−/−* cortices show a normal number of mitotic cells (PH3+, magenta) at the apical membrane but an increased number of mitotic cells basally. **(I)** The mitotic index of NSCs (Pax6+) is normal, but of BPs (Tbr2+) is significantly increased in *Cep55 −/−* cortices. Dashed line in A,D,G = apical membrane. For H,I: n= 4 *Cep55+/−* and 4 *−/−* brains. Scale bar: (A,G): 20 μm. n.s.; not significant, * p < 0.05; ** p < 0.01; *** p < 0.001. All experiments, Student’s t-test.

The neurons of the cp are daughters of cell divisions of NSCs and basal progenitors. NSCs divide many times, first symmetrically to multiply the NSCs, and then asymmetrically to produce BPs and neurons. BPs divide symmetrically to produce two neurons. Neuron daughters exit the cell cycle and differentiate. To determine whether the organization and numbers of NSCs and BPs in *Cep55* −/− cortices are altered, we used Pax6 and Tbr2 antibodies to mark them respectively **(Figure 2D)**. We observe a decrease in the number of NSCs in *Cep55 −/−* cortices, while the number of basal progenitors is normal **(Figure 2E)**. Notably, NSC and BP positioning appears disorganized, with a small but significant number of NSC nuclei mislocalized at positions basal to the svz **(Figure 2D – arrows; Figure 2E, F)**.

Cep55 is not thought to have a primary role in mitosis (Fabbro et al., 2005). To check this, we measured mitotic index in knockout brains, using phospho-histone3 (PH3) **(Figure 2G)**. There appears to be a trend for increased numbers of mitotic cells in *Cep55* −/− cortical sections (6.69 versus 9 cells per 100 μm; p = 0.11) (**Figure 2H)**; however, the increase is in basally rather than apically positioned mitotic nuclei, and is due to an increase in the mitotic index of BPs, not NSCs **(Figure 2I)**. This suggests there is not a primary defect in mitosis duration in *Cep55 −/−* NSCs. Overall, these data demonstrate there are reduced numbers of both NSCs and neurons in the *Cep55 −/−* cortex, with slight disorganization of layering, but not a significant increase in mitotic NSCs.

### Cep55 is expressed in proliferative cells in the brain and body, and is specifically detected in midbodies during late abscission

Having found defects in *Cep55 −/−* neurons, NSCs and BPs, we needed to ascertain in which of these cell types Cep55 is expressed. Previous RNA sequencing experiments found *Cep55* mRNA expressed in proliferating cell types in early mouse embryos (Gao et al., 2017); and in the murine cerebral cortex at E14.5 and P0 (Loo et al., 2019). RNA *in situ* hybridization shows *Cep55* is expressed in proliferative zones of the cortex and eye at E14.5, but is not detectable in neuronal layers. Lower expression in other proliferating tissues is observed **(Figure 3 – figure supplement 1A-C)**. We tested for endogenous Cep55 protein expression and subcellular localization by immunohistochemistry on control and knockout cells, and detected Cep55 protein in dividing NSCs of control embryos, but not knockouts **(Figure 3A-C)**. Interestingly, Cep55 is specifically detected only at late stages of abscission: in late midbodies as a disk at the center of the midbody, and in the post-abscission midbody remnants (MBRs). No specific Cep55 signal was detected in metaphase, anaphase, or early midbodies. In basal progenitors from control brains, Cep55 has the same localization pattern **(Figure 3D)**. We do not detect Cep55 protein at centrosomes **(Figure 3E)**, concurring with others (Bastos & Barr, 2010; Zhao et al., 2006).

**Figure 3:**
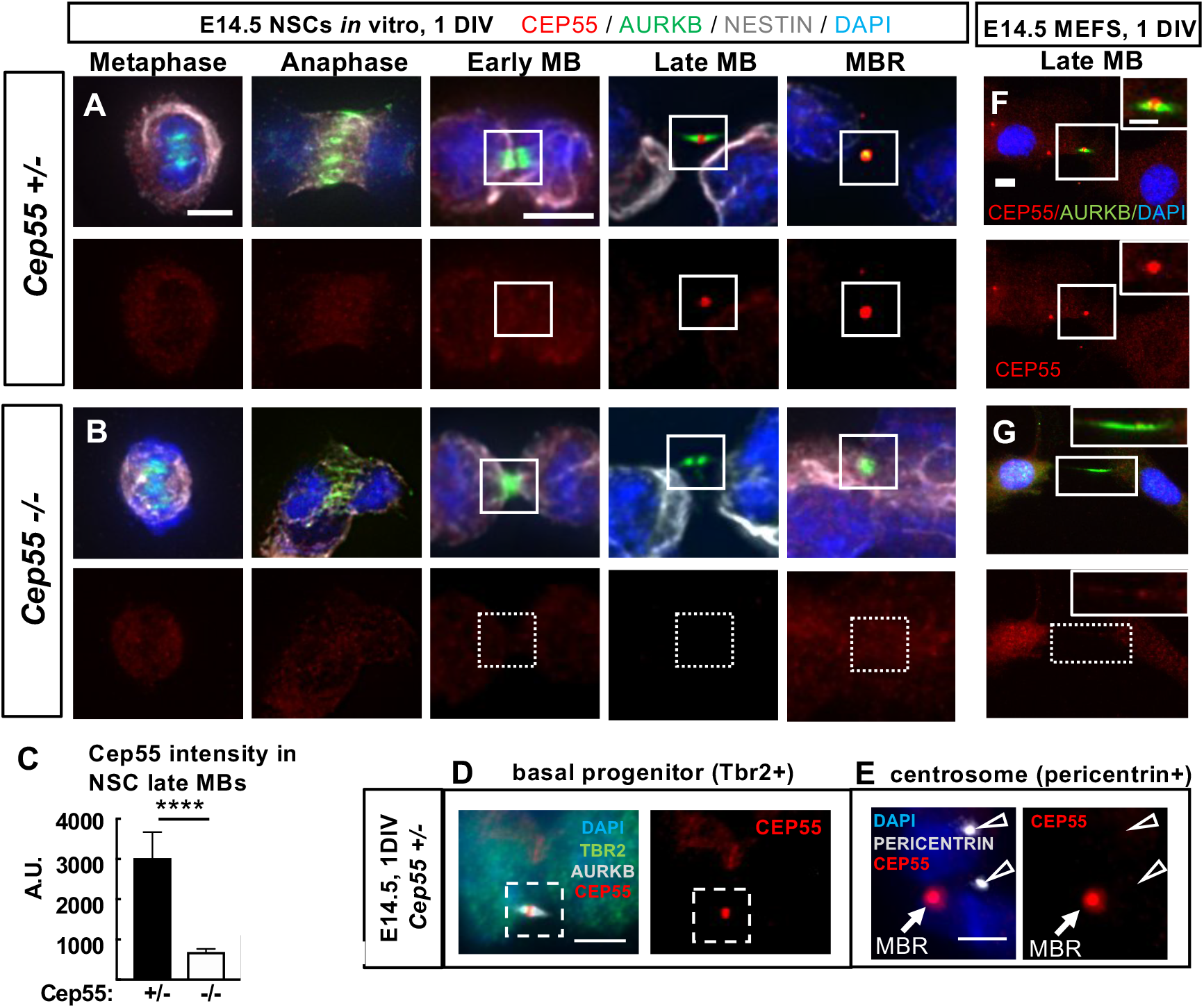
Cep55 protein localizes in late-stage midbodies of NSCs, basal progenitors and MEFs. **(A, B**) Representative images of control (*+/−*) and *Cep55* knockout (*−/−*) E14.5 neural stem cells (NSCs) undergoing cytokinesis that were cultured for 1 day in vitro (DIV) fixed and immunostained for endogenous Cep55 (red), Aurora kinase B (green), Nestin for NSCs (grey) and DAPI. Cep55 is not detectable in mitotic spindles or centrosomes in metaphase, anaphase, or early midbody stage NSCs, but does accumulate in a disk in the center of late midbodies and in midbody remnants in control (A) but is undetectable in knockout (B) NSCs. **(C)** Quantification of endogenous Cep55 IF signal intensity in control and *−/−* NSCs midbodies. **(D)** Cep55 in a late-stage midbody of a basal progenitor from control brain. **(E)** Cep55 signal was not detected at centrosomes, marked by pericentrin, in NSCs (open arrows), but is detected in a nearby midbody remnant (closed arrow). **(F, G)** *Cep55+/+* MEFs show a similar pattern of Cep55 localization as NSCs, with accumulation only in late-stage midbodies (F), but is undetectable in *Cep55−/−* MEF midbodies (G). Scale bars: A, metaphase and early midbody: 5 μm; D: 4 μm; E: 5 μm; F: 10 μm, inset 5 μm. ****, p <0.0001, Student’s t-test.

To investigate whether Cep55 is expressed in cells of non-neural tissues, we dissociated mouse embryonic fibroblasts (MEFs) from E14.5 control and *Cep55* −/− embryo bodies. We noted the same expression pattern in MEFs as in NSCs, with detectable signal above background only in late-stage midbodies of control MEFs **(Figure 3F, G)**. Thus, we find Cep55 is expressed in multiple proliferating embryonic cell types – NSCs, BPs and MEFs – and that it accumulates specifically at the late stage of abscission in the central bulge of the midbody.

### Midbody defects in fixed cortices of *Cep55* knockouts are consistent with delays in NSC abscission

Based on the specific localization of Cep55 to late-stage NSC midbodies, as well as the prior reports of a requirement for Cep55 in completion of abscission in cell lines, we hypothesized that microcephaly in *Cep55* −/− mice is due to a primary defect in abscission completion. However, the growth of the brain (albeit less than normal), and almost normal body size of *Cep55* knockouts suggest that abscission is completing in most cells. To begin to address whether abscission processes in *Cep55* −/− brains are normal, we used a fixed cortical slab preparation in which we can image and quantitatively analyze many NSCs undergoing mitosis and cytokinesis at the ventricular surface **(Figure 4A-B)** (Janisch & Dwyer, 2016; Janisch et al., 2013). By labeling the NSC apical cell junctions with ZO-1, we observe that *Cep55* −/− endfeet are more variable in size, with many abnormally large endfeet (**Figure 4C)**. Indeed, quantification shows a striking decrease in the density of apical endfeet **(Figure 4D)**. Since the apical membrane expands during mitosis, we quantified the apical mitotic index of NSCs, using phospho-histone H3 (PH3), but it is not altered in *Cep55* −/− cortices **(Figure 4C - left panels, E)**, consistent with our finding in cross sections **(Figure 2H)**. However, the midbody index, the percent of NSCs in abscission, is increased by approximately 25% at E14.5 **(Figure 4C - right panels, F)**. These data suggest that in *Cep55* knockout cortices, NSCs may take longer to complete abscission. As the abscission process proceeds, midbodies mature by becoming thinner, and then forming constriction sites on each flank, where microtubule disassembly and membrane scission occurs (Guizetti et al., 2011). We observe fewer midbodies with constriction sites in *Cep55* −/− brains **(Figure 4G),** and more short midbodies **(Figure 4H).** There was no significant difference in midbody widths (data not shown). These data suggest that without Cep55, midbodies are able to compact their microtubules to become thinner as they mature, but have a defect in making the constriction sites, and abscission duration is longer than normal.

**Figure 4:**
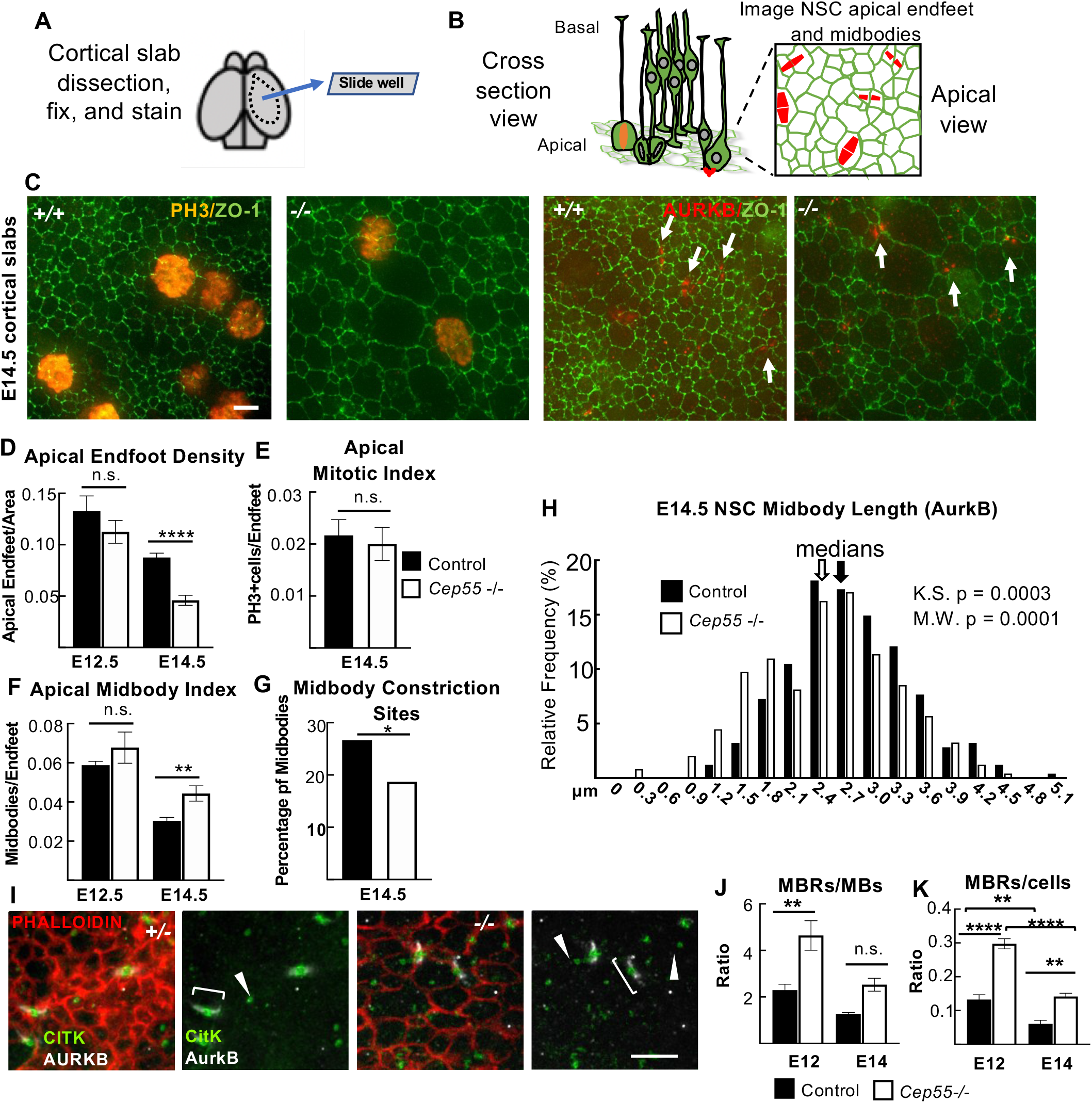
*Cep55* knockouts display NSC midbody defects in cortical slab preparations. **(A, B)** Schematics of cortical slab dissection, and cross section vs. apical membrane views of NSCs undergoing cytokinesis. **(C)** E14.5 cortical slabs immunostained for apical junctions (ZO-1, Zona-Occludens-1), and mitotic chromatin (PH3+) or midbodies (AURKB, Aurora B kinase). **(D)** Mean apical endfoot density is reduced in *Cep55 −/−* at E14.5. For D, E12.5: N= 4 *+/−* slabs (4 brains), 4 *−/−* slabs (4 brains). E14.5: N = 8 control slabs (6*+/+*, 2*+/−* brains) and *7 −/−* slabs (6 brains). (**E-F)** Mitotic index is normal, but midbody index is significantly increased at E14.5. For E, N= 6 control slabs (4*+/+*, 2*+/−* brains), 6 *−/−* slabs (5 brains). For F, E12.5: N= 4 *+/−* slabs (4 brains), 4 *−/−* slabs (4 brains). E14.5: N= 6 control slabs (4 *+/+*, 2*+/−* brains), 5 *−/−* slabs (5 brains). **(G)** A smaller percentage of Cep55 *−/−* midbodies have visible constriction sites. **(H)** Cep55 *−/−* midbodies tend to be shorter than controls (measured by AurkB immunolabeling). Medians: 2.7 μm for +/+, 2.5 μm for *−/−*. Bin = 0.3μm. For G, H, N= 353 control midbodies (5*+/+*, 2*+/−* brains), 246 *−/−* midbodies (5 brains) **(I)** E14.5 slabs immunostained for AurkB to mark pre-abscission midbody flanks (brackets), and citron kinase (CitK) to mark midbody bulges and post-abscission MBRs (arrowheads). **(J,K)** MBRs are increased in *Cep55−/−* brains, normalized to MB number or NSC (endfoot) number. For J,K, E12.5 and E14.5: N= 4 control slabs (2 *+/+*, 2*+/−* brains), 4 *−/−* slabs (4 brains). Scale bars: B: 2 μm I: 5μm. *p<0.05, **p<0.01, ***p<0.001, ****p<0.0001, n.s., not significant. D,F-G: Student’s T-test, J,K: ANOVA, E: Fisher’s exact test.

Since Cep55 protein is abundant in midbody remnants (MBRs) **(Figure 3A)**, we investigated whether Cep55 loss alters MBR numbers at the apical membranes of NSCs, using Citron kinase (CitK) as a marker (Ettinger et al., 2011; Gruneberg et al., 2006). Remarkably, there are approximately twice as many MBRs present on the apical membranes of *Cep55 −/−* brains as in control brains, whether normalized to midbody number or cell number **(Figure 4I-K)**. This could be another manifestation of delayed abscission, at a late stage when the flanks are no longer detectable with Aurora B staining. It could also represent a role of Cep55 in MBR disposal. Altogether, theses analyses in fixed brains show midbody abnormalities that are consistent with delayed abscission of NSCs when Cep55 is absent.

### Cep55 knockout NSCs can complete microtubule disassembly, but it is delayed

To directly test whether abscission is delayed or fails in NSCs of *Cep55 −/−* cortex, we performed time-lapse imaging of abscission *in vivo*, in live cortical slab explants, using a method we developed previously (McNeely & Dwyer, 2020). By dissecting cortical slabs from embryonic brains of a membrane-GFP mouse line, and incubating them with the cell-permeable far-red fluorescent microtubule dye SiR-tubulin, we can image individual NSCs at the apical membrane undergoing cytokinesis from furrowing to abscission. As the cleavage furrow ingresses towards the apical membrane, the midbody forms as a tight bundle of microtubules, the central bulge becomes apparent, and then microtubule disassembly occurs on each midbody flank independently **(Figure 5A)**. We previously showed that in control E13.5 NSCs *in vivo*, abscission takes on average 50 minutes to complete, with a wide range of times observed, and microtubule disassembly occurs on both sides of the midbody flank in the majority of divisions (McNeely & Dwyer, 2020).

**Figure 5:**
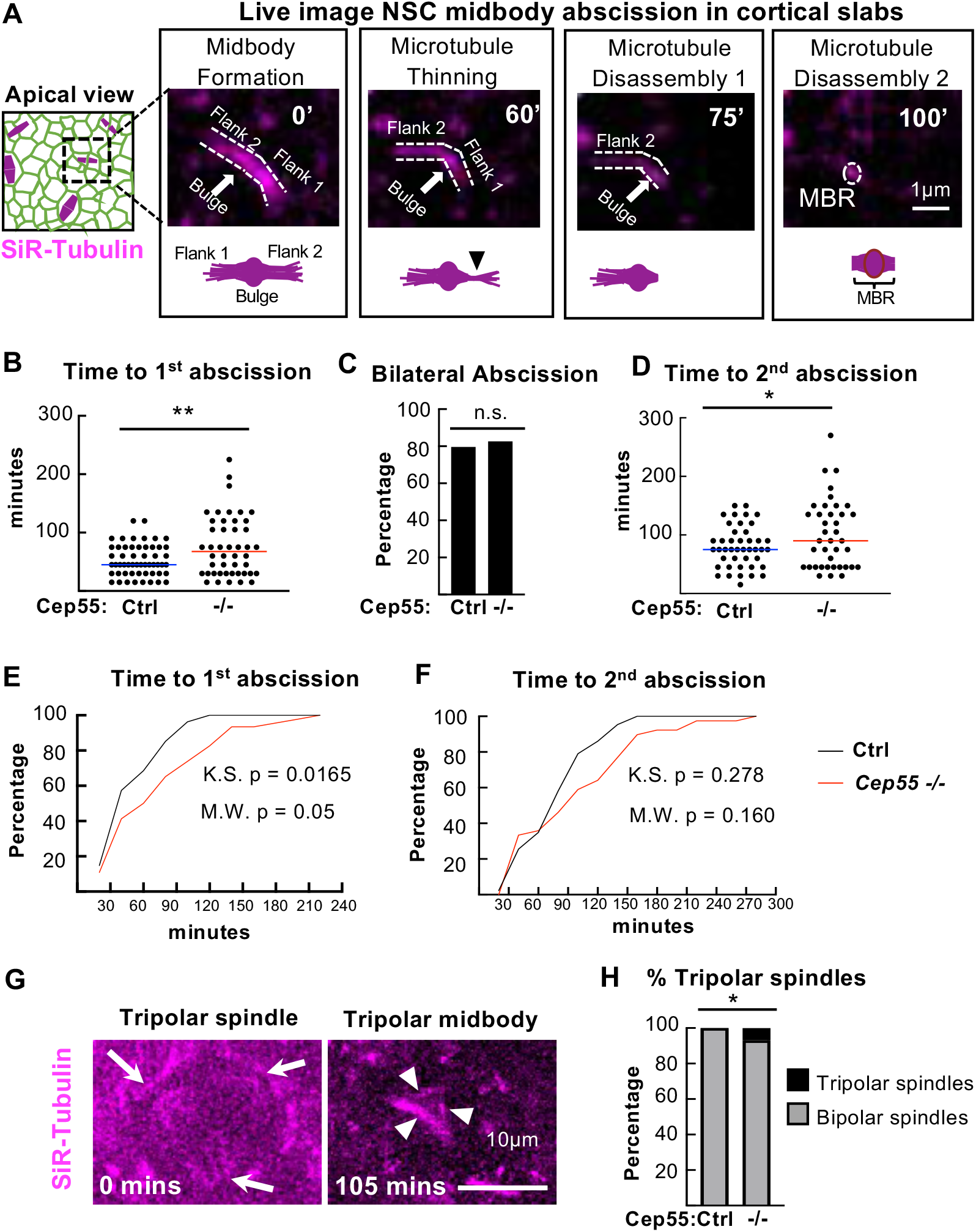
*Cep55* knockout causes delayed midbody microtubule disassembly during NSC abscission. **(A)** Schematics and time-lapse images of an E13.5 NSC in a cortical slab explant undergoing midbody abscission. Microtubules are labeled by SiR-Tubulin. Distinct steps shown: midbody formation, flank 1 thinning, microtubule disassembly on flank 1 (1st abscission), and microtubule disassembly on flank 2 (2nd abscission). After bilateral flank disassembly, the midbody remnant (MBR) is left at the apical membrane. **(B-D)** Time from midbody formation to complete 1^st^ abscission and 2^nd^ abscission are increased in *Cep55−/−* NSCs, but there is no change in the percentage of bilateral abscissions detected. **(E, F)** Cumulative frequency plots for the 1^st^ abscission and 2^nd^ abscission show the curves are shifted to the right in *−/−* NSCs. **(G)** *Cep55−/−* NSC with a tripolar spindle (arrows point to spindle poles) proceeded to form a tripolar midbody at 105 minutes. (arrowheads point to midbody “flanks”). **(H)** Tripolar spindles are increased in *Cep55* −/− cortices. For B,C, E, N= 54 control cells (2 *+/+*, 1 *+/−* slabs), 46 *−/−* cells (4 slabs); D,F: N= 43 control cells,39 *−/−* cells. G: N= 71 control cells (2*+/+*, 1*+/−* slabs) and 86 *−/−* cells (4 slabs). *p<0.05, **p<0.01. B,D: T-test. E,F: K-S and M-W tests. C,H: Fisher’s Exact test.

Using this method, we quantified several aspects of the abscission process in *Cep55 −/−* NSCs: midbody formation, midbody structure, and time to abscission (ascertained as microtubule disassembly) on one or both flanks. Unexpectedly, we find the vast majority (92%) of *Cep55* −/− NSCs are able to complete abscission. However, these knockout NSCs do have a significant increase in average time to the first abscission (delay of 23 minutes, **Figure 5B**). They are also able to complete bilateral abscissions: eighty percent of both control and *Cep55* −/− NSCs have observable second abscissions on the other midbody flank **(Figure 5C)**. But again, the time to the second abscission is significantly increased in knockout NSCs **(Figure 5D)**. Cumulative frequency plots illustrate the slower abscission kinetics in the *Cep55* −/− NSCs **(Figure 5E-F)**.

We did observe a small number of *Cep55* knockout NSCs that displayed abnormalities at an earlier step of cytokinesis. Notably, there is a significant increase in the number of *Cep55 −/−* NSCs that have tripolar mitotic spindles (7%, 6 out of 81 knockout NSCs imaged versus 0 out of 71 in controls) **(Figure 5G-H)**. Of these 6 NSCs with tripolar spindles, four progressed into tripolar midbodies, which remarkably were still able to undergo microtubule disassembly, one regressed, and one drifted out of the imaging field. We observed a few *Cep55 −/−* NSCs that had bipolar spindles that initiated a furrow but did not complete (4/50, compared to 0/54 control NSCs). In three of these NSCs, the furrow only partly ingressed then regressed, so the midbody never formed, while in the other, the midbody formed, but then the membrane regressed. Taken together with our preceding data on Cep55 protein localization and fixed midbody abnormalities, the findings indicate that the primary function of Cep55 in NSCs is to promote timely abscission, with particular effects on the kinetics of late steps such as constriction site formation and microtubule disassembly.

### *Cep55* knockout NSCs and MEFs have decreased but not eliminated ESCRT recruitment to late-stage midbodies

Since *Cep55 −/−* NSCs have delayed abscission, and Cep55 is thought to recruit other abscission proteins, we wanted to determine if other known abscission proteins were altered. The current model for the mechanism of abscission, developed from mammalian cell lines and invertebrates, proposes that at the late stage of abscission, endosomal sorting complexes required for transport (ESCRT) components are recruited to the midbody, and then form helical filaments extending from the central bulge to constriction sites, compacting microtubules and pulling the plasma membrane close enough for membrane scission (Christ et al., 2017; Goliand et al., 2018; Guizetti et al., 2011; Stoten & Carlton, 2018). Cep55 is thought to be necessary to recruit this abscission machinery to the midbody in human cells through interactions with ESCRT-I/TSG101 and Alix. Then TSG101 and Alix both recruit ESCRT-III, which assemble into filaments (Carlton et al., 2008; Christ et al., 2016; Lee et al., 2008; Morita et al., 2007) **(Figure 6A)**. Notably, Cep55 is absent in invertebrate genomes, yet their midbodies still recruit ESCRTs to mediate abscission. Therefore, we asked whether abscission in *Cep55 −/−* NSCs is accomplished with or without the ESCRT machinery. First, we tested for the localization of endogenous Tsg101 and Alix in midbodies of control and *Cep55 −/−* NSCs. Approximately one-third of control NSC midbodies have Alix or Tsg101 present at the midbody center, reflecting that recruitment is temporally regulated and occurs only at late abscission stages. Among *Cep55 −/−* NSC midbodies, however, only about 5% had detectable Alix and Tsg101 (**Figure 6B)**. This suggests that in the absence of Cep55, ESCRT component recruitment may be delayed, or total protein recruitment may be decreased.

**Figure 6:**
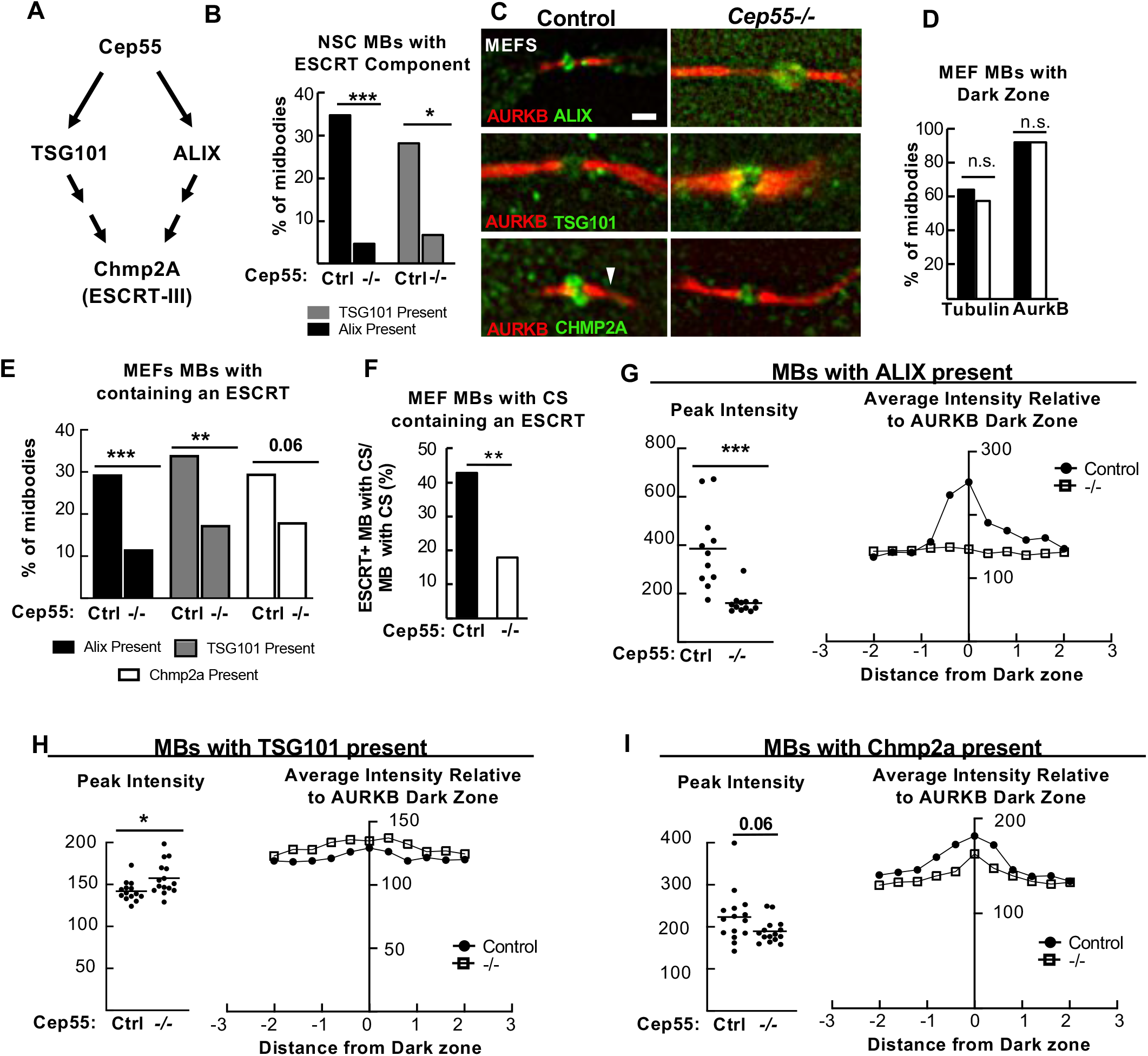
*Cep55* knockout NSCs and MEFs have reduced but not eliminated ESCRT recruitment to midbodies. **(A)** Model from literature for Cep55 recruiting ESCRTs in midbodies (MBs). **(B)** *Cep55−/−* NSC midbodies are much less likely to have detectable Alix or TSG101. Alix: N= 49 NSC control MBs (3 animals, 3 coverslips); N= 46 *−/−* MBs (2 animals, 3 coverslips); TSG101: N= 32 control (2 animals, 2 coverslips), N= 45 *−/−* (3 animals, 3 coverslips). All E14.5 NSCs 1 DIV. **(C)** Representative images of MBs of Mouse embryonic fibroblasts (MEFs) cultured for 1 day, from control or *Cep55−/−* E14.5 embryos, immunostained for ALIX, TSG101, or Chmp2A. Arrowhead, constriction site. Scale Bar 1μm. **(D)** Central dark zones were detected at normal percentage in *−/−* MEF MBs, using alpha-tubulin or AurkB immunostaining. For D: N= 402 control, 355 *−/−* MBs (5 animals, 6 coverslips each). **(E)** *Cep55−/−* MEF midbodies are much less likely to have detectable Alix, TSG101, or Chmp2a than controls. Alix: N= 153 control, 129 *−/−* MBs (4 animals, 4 coverslips each); Tsg101: N= 141 control, 132 *−/−* MBs (4 animals, 4 coverslips each); Chmp2a: N= 108 control, 105 *−/−* MBs (3 animals, 3 coverslips each). **(F)** The percentage of MBs with constriction sites (CS) that contain detectable ESCRT is decreased in *−/−* MBs. N (midbodies with constriction sites) = 110 control, 95 *−/−*, (5 animals, 11 coverslips each). **(G-I)** MBs with ESCRT present were analyzed by fluorescence intensity linescans through the central dark zones. Alix peak is severely reduced in *−/−* MBs, but Tsg101 and Chmp2a peaks are not. Alix: N=11 control,*−/−* MBs (2 animals, 2 coverslips, 1 experiment); Tsg101: N=14 control, 15 *−/−*MBs (2 animals, 2 coverslips, 1 experiment); Chmp2a: N= 15 control, *−/−* MBs (2 animals, 2 coverslips, 1 experiment). * p < 0.05; ** p < 0.01; *** p < 0.001. C-G: Fisher’s exact test; H-J,: Student’s t-test.

Next, we used MEFs to further analyze midbodies and ESCRT recruitment when Cep55 is absent, since MEF cells and midbodies are much flatter than NSCs and easier to image. Control MEFs exhibit identical Cep55 localization in late-stage midbodies as NSCs do **(Figure 3F)**. Additionally, an antibody against the downstream ESCRT-III component, Chmp2a, works on MEFs **(Figure 6C)**. Cep55 was previously suggested to be required for formation of the dense central midbody matrix in HeLa cells, which appears as an unlabeled “dark zone” with alpha-tubulin or Aurora B antibody immunostaining (Zhao et al., 2006). We tested this in MEFs, but the dark zones were equally detected in *Cep55* −/− MEFs as controls **(Figure 6D)**. *Cep55 −/−* MEF midbodies had similar length and width to controls (data not shown). Next, we tested Alix and Tsg101 recruitment, and found similarly to NSCs, the percentages of *Cep55* −/− MEF midbodies with detectable endogenous Alix and TSG101 are significantly decreased compared to controls **(Figure 6C, E)**. There is also a trend for reduced midbodies with the ESCRT-III component Chmp2a detectable **(Figure 6E)**. Interestingly, if we just consider midbodies that have formed constriction sites, about 40% of control MEFs have ESCRT localization, versus 15% of the *Cep55 −/−* MEFs, suggesting that constriction sites can form before or independently of ESCRT recruitment **(Figure 6F).**

Next, we asked whether the minority of *Cep55 −/−* MEF midbodies with detectable ESCRTs had normal distribution of that ESCRT within the midbody. We evaluated ESCRT protein signal intensity using line scans drawn lengthwise along the midbody. In controls, a peak of ESCRT intensity is seen as expected in the central matrix/ “dark zone” between the Aurora B stained flanks **(Figure 6G-I, controls)**. In *Cep55 −/−* midbodies that had detectable Alix, the average peak intensity of Alix fluorescence is significantly decreased, and the peak of intensity at the dark zone is lost **(Figure 6G)**. Surprisingly, for Tsg101-containing *Cep55 −/−* midbodies, there is not a decrease but a slight increase in peak intensity of Tsg101, and the intensity pattern is similar to controls **(Figure 6H)**. We found a trend for decreased peak Chmp2a fluorescence compared to controls **(Figure 6I)**. This suggests the interesting possibility that Alix and Tsg101 recruitment are affected differently by loss of Cep55. Together these data indicate that Cep55 is not essential for ESCRT component recruitment to the midbody in NSCs and MEFs, but suggests that Cep55 may increase or accelerate their recruitment to ensure efficient abscission.

### *Cep55* knockout mice have increased numbers of binucleate cortical cells and MEFs

Our analyses of midbodies and abscission in NSCs of the *Cep55* knockout so far suggest that most NSCs can complete abscission in the absence of *Cep55*, but that the process is slower and a subset of cells may fail. To assay cytokinesis success and failure in a larger sample of cells, we analyzed DNA content of embryonic cortical cells with flow cytometry. Dissociating cells of E15.5 cortices and labeling DNA with propidium iodide, we found 30% of *Cep55 −/−* cortical cells have 4N DNA content compared to 2% in controls, a 15-fold increase **(Figure 7A)**. An increase in cells with 4N DNA content could come from either an arrest in G2 /M phase, or failed cytokinesis resulting in formation of a binucleate or tetraploid progenitor or neuron. Since we did not see significantly increased numbers of PH3+ cells in *Cep55 −/−* cortices **(Figure 2H)**, this suggests the presence of binucleate/tetraploid cells. To investigate whether cells with 4N DNA content were progenitors or neurons, we used Ki67 and DAPI co-labeling to differentiate cycling progenitor cells (Ki67+) from non-cycling neurons (Ki67-) cells. Interestingly, we observed an increased 4N DNA peak in both Ki67+ **(Figure 7B)** and Ki67-*Cep55* −/− cells **(Figure 7C)**. In controls, there is no 4N DNA peak in Ki67-negative cells, because neurons are post-mitotic **(Figure 7C, left)**. These data suggest that in *Cep55* −/− cortices, there are significant numbers of binucleate neurons and NSCs.

**Figure 7:**
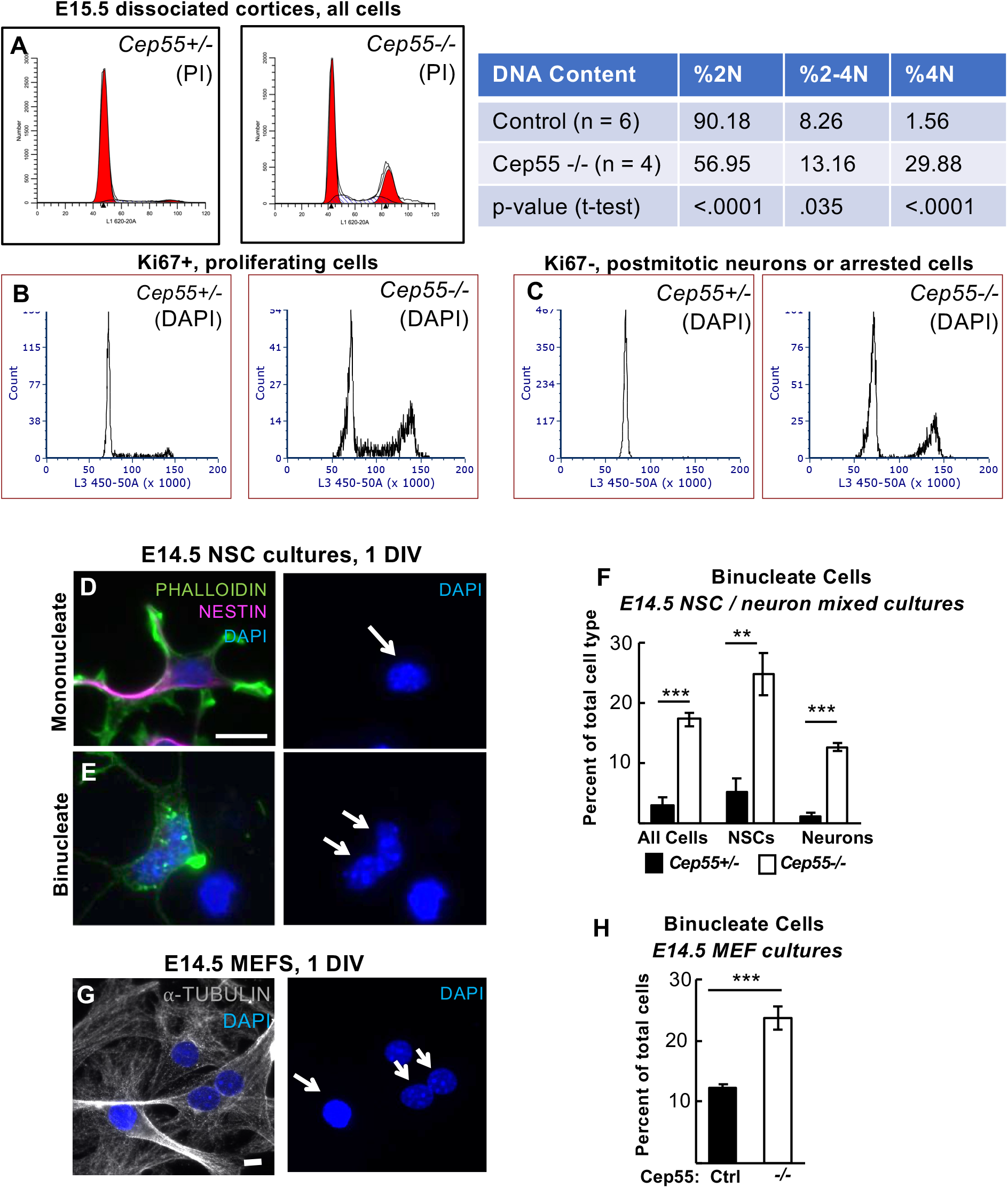
*Cep55* knockout mice have increased numbers of binucleate cortical cells and MEFs. **(A)** Flow cytometric analysis of E15.5 dissociated cortices labeled with propidium iodide (PI) indicates an increase in cells with tetraploid (4N) DNA content in *Cep55−/−* brains, and concomitant decrease in cells with 2N DNA content. **(B-C)** Both proliferating (Ki67+) and non-proliferating (Ki67-, presumed neurons) populations of *Cep55−/−* brains show increases in 4N DNA cells. **(D-F) I** ncreased numbers of binucleate progenitors (Nestin+) and neurons (Nestin-) are seen in *Cep55−/−* dissociated cortical cultures at 1 day in vitro (DIV). D, arrow: example of mononucleate progenitor. E, arrow: example of binucleate neuron. **(G-H)** Primary cultures of mouse embryonic fibroblasts (MEFs) from *Cep55−/−* embryos have twice as many binucleate cells as control cultures at 1DIV. Arrow: example of mononucleate cell, double arrow: example of binucleate cell. For (A) n= 6 *Cep55+/+*;*+/−*, 4 *Cep55−/−* dissociated cortices; (B-C) n= 5 *Cep55+/+;+/−*, 5 *Cep55−/−* dissociated cortices; (F) n= 4 *Cep55+/+;+/−* and 4 *Cep55−/−* coverslips from 2 embryos each. (H) n=5 *Cep55+/−,* 5 *Cep55−/−* coverslips from 3 embryos each. n.s.; ** p < 0.01; *** p < 0.001. Scale bars in D and G, 10 μm.

To further investigate whether there are binucleate NSCs and/or neuron populations in *Cep55 −/−* cortices, we imaged dissociated, cultured cortical cells from E14.5 cortices. We used Nestin immunostaining to mark NSCs, and actin (phalloidin) and DAPI staining to differentiate cells with one or two nuclei. Indeed, we observed increased percentages of binucleate cells in *Cep55 −/−* cultures, including approximately 24% of NSCs (Nestin+) and 12% of neurons (Nestin-) **(Figure 7D-F)**. These data show that *Cep55* knockout results in significant numbers of binucleate NSCs and neurons in the brain.

Since the brain size is more severely affected than body size in *Cep55* knockouts **(Figure 1D)**, We wondered whether binucleation is a consequence of *Cep55* loss in non-neural cell types too. Cep55 is localized in embryonic fibroblast midbodies in a similar distribution as seen in NSCs **(Figure 3F)**, and its loss causes decreased recruitment of ESCRTs **(Figure 6E)**. Indeed, we do observe a doubling of binucleate cells in *Cep55* −/− MEF cultures compared to controls **(Figure 7G, H)**. Thus, while not as dramatic as seen in NSC cultures, Cep55 loss does lead to binucleation in MEFs as well.

### Apoptotic cells are increased in *Cep55* knockout neural but not non-neural tissues during embryogenesis

Abscission defects and binucleation have been linked to apoptosis of NSCs and neurons in other genetic mouse microcephaly mutants (Bianchi et al., 2017; Little & Dwyer, 2019; Moawia et al., 2017). The disorganization and reduced cell numbers we noted in *Cep55 −/−* cortices suggest some cells could be dying. To assay for apoptosis, we labeled cortical sections with antibodies to cleaved-caspase 3 (CC3). In control developing cortex, apoptotic cells are only rarely detected, but there is a striking increase in apoptotic cells in *Cep55 −/−* cortices **(Figure 8A)**. Apoptosis is most increased in the proliferative zones, but is also increased in the cortical plate (cp, neuronal layer) **(Figure 8B)**. This suggests widespread apoptosis in multiple cortical cell types, with the highest increase in dividing cell types, NSCs and BPs. Since brain size is already significantly decreased at E14.5, we looked at an earlier age for apoptosis. Indeed, apoptosis is dramatically increased in *Cep55* −/− cortical epithelium at E10.5, when the cortex consists only of NSCs **(Figure 8C-D)**. Therefore, we conclude that apoptosis of primarily NSCs but also neurons contributes to impaired cortical growth.

**Figure 8:**
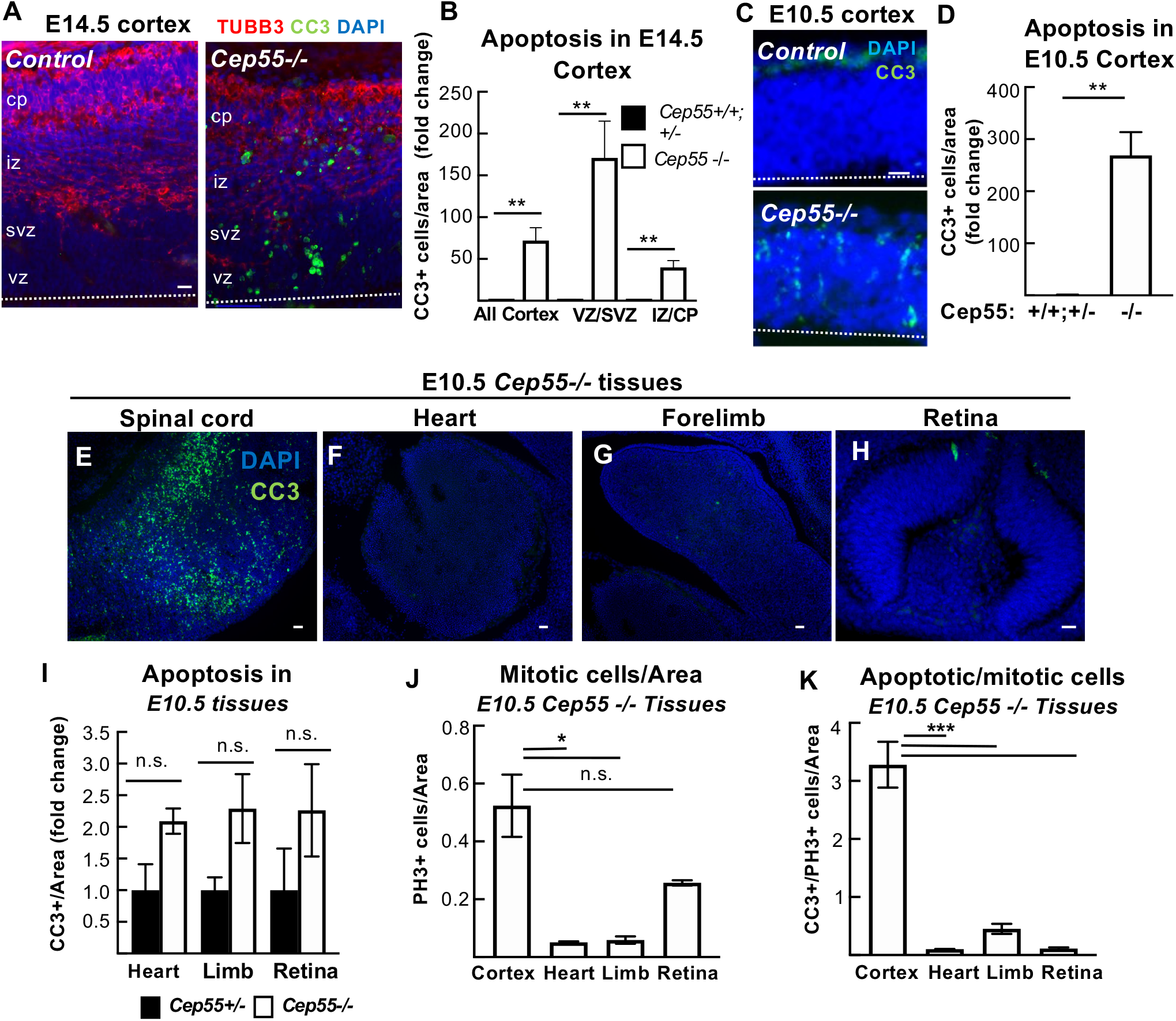
Apoptosis is greatly increased in the neural tissue of *Cep55* knockout embryos. **(A)** Cortical sections immunostained for apoptotic cell marker cleaved-caspase 3 (CC3, green) show almost no CC3+ cells in control but many in *Cep55−/−* cortex. **(B)** At E14.5, apoptotic cells were most increased in the vz/svz, containing nuclei of NSCs and BPs, but also increased in neurons in cp/iz of *−/−* brains. **(C-D)** At E10.5, prior to neurogenesis, apoptosis is greatly increased in *−/−* forebrain NSCs.**(E-I)** Apoptosis is high in the E10.5 *−/−* spinal cord, (and midbrain and hindbrain, not shown), but not in heart, forelimb, or retina. **(J-K)** The brain specificity of the apoptosis is not simply due to higher mitotic index. Dashed line in A,C = apical membrane. For all experiments, n = 3 control and −/− brains or embryos at each age. Scale bars: (A): 20 μm (C): 20 μm, (D): 40 μm, (E): 40 μm, (I) 20 μm. * p <0.05, ** p < 0.01, *** p < 0.001. n.s., not significant. All experiments, Student’s t-test.

We wondered whether a lack of apoptosis in non-cortical tissues could explain the less severe body size in *Cep55 −/−* mice compared to brain size. Alternatively, apoptosis could occur in these tissues, but proliferation increases to compensate. To address this question, we labeled E10.5 whole-embryo sections with CC3. Interestingly, apoptosis is observed throughout central nervous system tissues **(Figure 8E**, spinal cord, and hindbrain and midbrain, not shown) but is not seen in the rest of the body, including the heart precursor **(Figure 8F)**, the forelimb **(Figure 8G)**, and the retina **(Figure 8H)**; quantification **(Figure 8I)**. There are many more mitotic cells in the developing cortex than in these tissues in both controls (data not shown) and *Cep55 −/−* mice **(Figure 8J)**; however, even when normalized to the number of mitotic cells, apoptosis is specifically increased in the *Cep55 −/−* cortex, not in the other tissues **(Figure 8K)**. These data suggest the interesting possibility that a distinct apoptotic response occurs in central nervous system tissues after *Cep55* loss, that does not occur in all proliferating tissues.

### p53 nuclear expression is increased in *Cep55* knockout binucleate cortical cells, but not in binucleate MEFs

We previously showed that NSC apoptosis in a different abscission mutant, in the kinesin Kif20b, was mediated by the tumor suppressor p53 (Little & Dwyer, 2019). To determine whether p53 elevation occurs in *Cep55 −/−* cortices, we labeled E14.5 cortical sections with antibodies to p53. Indeed, while virtually no cells with bright nuclear p53 accumulation are observed in control sections, greatly increased numbers of p53+ cells are seen in *Cep55 −/−* sections **(Figure 9A-B)**. We noticed these cells throughout the cortex, but especially increased in proliferative zones (vz/svz). While some p53 positive cells appear singular, others appear paired **(Figure 9A, arrows)**. To further delineate in which *Cep55 −/−* cells p53 expression occurred, we used dissociated cortical cell cultures. We observed a 5-fold increase in the number of cells with a nuclear:cytoplasmic (N:C) ratio of 2 or greater, indicative of p53 activation as it acts in the nucleus **(Figure 9C-D)**.

**Figure 9.**
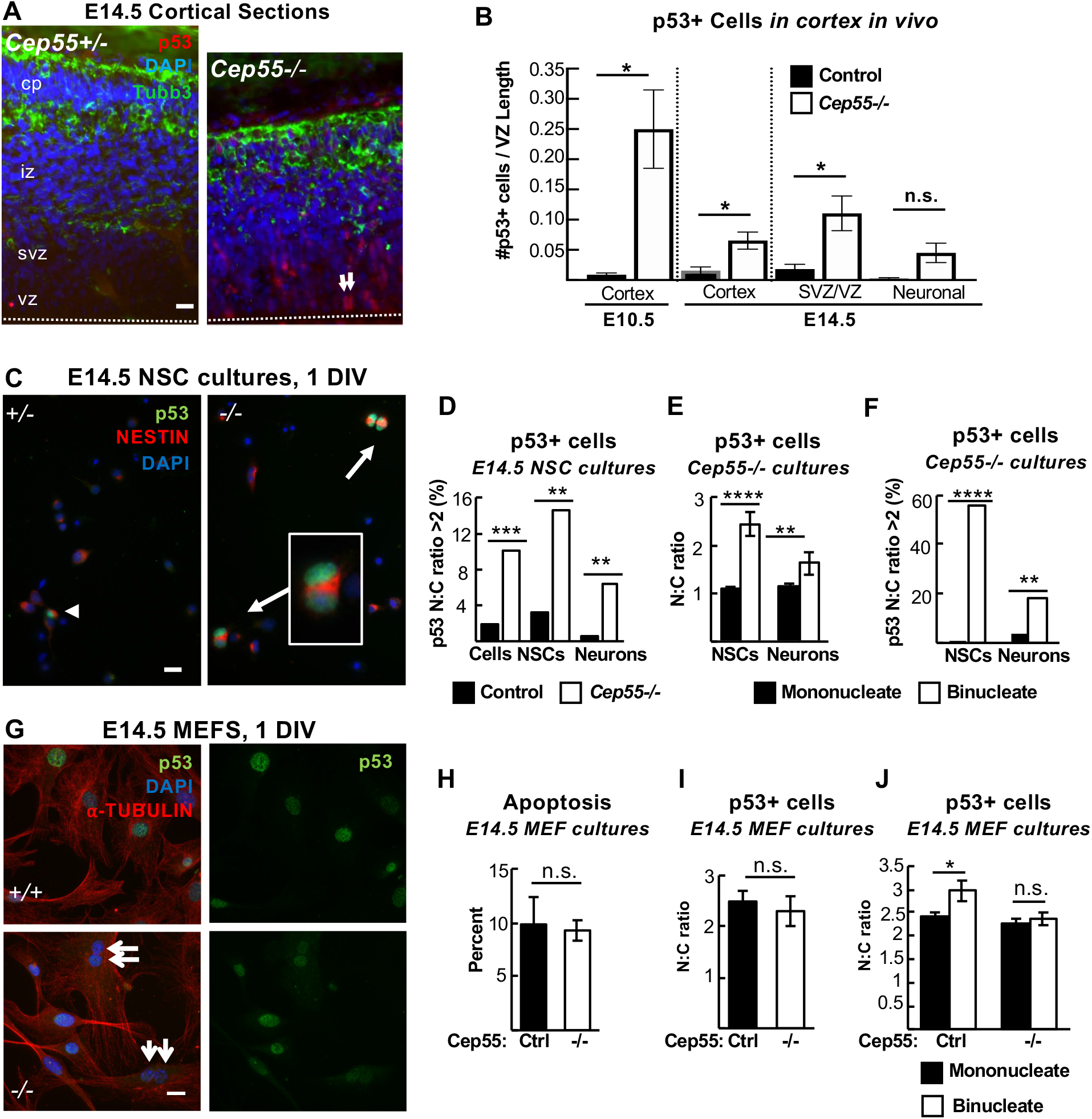
p53 nuclear accumulation is increased in *Cep55* knockout binucleate NSCs and neurons, but not in binucleate MEFs. **(A-B)** Cortical sections immunostained for p53 and Tubb3 show almost no p53+ cells in controls but many in *Cep55−/−* cortices. Arrows, paired nuclei with p53 expression. **B.** p53+ cell counts are increased in *Cep55−/−* cortices at E10.5, and at E14.5, mostly in the vz/svz than neuronal layers **(C)** Images of dissociated NSC cultures show p53+ binucleate NSCs from *Cep55 −/−* cortices. **(D)** Counts of cells with a p53 nuclear:cytoplasmic (N:C) ratio of >2 are greatly increased in *−/−* cultures, both NSCs and neurons. **(E-F)** In Cep55 *−/−* NSC cultures, the mean N:C ratio of p53 intensity is about 1 in mononucleate NSCs or neurons, but significantly higher in binucleate progenitors and neurons. **(F)** In *Cep55−/−* cultures, over half of binucleate NSCs have a p53 N:C ratio > 2, compared to only 1% of mononucleate NSCs. Among binucleate neurons, ~ 20% have a p53 N:C ratio of >2 versus only 2% of mononucleate neurons. **(H)** Apoptosis (CC3+) is not increased in *Cep55−/−* primary MEF cultures compared to controls. **(I)** The N:C ratio of p53 signal (p53, green, G) is not different in *−/−* MEFs compared to controls. **(J)** Binucleate *Cep55 −/−* MEFs did not have an increased p53 N:C ratio compared to mononucleate *−/−* MEFs (G, arrows). Dashed line in A = apical membrane For B, n = 3 Cep55+/+;+/− and 3 Cep55 −/− mice at each age. For D-F, n= 4 Cep55+/+;+/− and 4 Cep55−/− coverslips from 2 embryos each; 124 control and 137 −/− NSCs; 143 −/− and 184 −/− neurons for N:C ratios. For H, n= 3 control and 3 Cep55−/− coverslips from 3 embryos each. For I, n= 195 control and 232 −/− cells from 3 control and Cep55−/− coverslips and embryos each. For J, n= 153 mononucleate and 25 binucleate control cells and 158 mononucleate and 68 binucleate −/− cells from 3 coverslips and 3 embryos each. Scale bars in A, C, G represent 20 μm. * p < 0.05; ** p < 0.01; *** p < 0.001; **** p < 0.0001, t-test.

Interestingly, just as apoptosis is increased in both NSCs and neurons in *Cep55 −/−* cortex, we found increased nuclear p53 expression in both cell types **(Figure 9D).** While cytokinetic defects would occur only in dividing NSCs, we reasoned that a failed cytokinesis event could result in the formation of a binucleate daughter cell, either progenitor cell or neuron. Indeed, we had observed both binucleate NSCs and neurons in *Cep55 −/−* cultures **(Figure 7F)**. To investigate if binucleation was associated with p53 activation and apoptosis in progenitor cells and neurons, we co-labeled cells with Nestin, Phalloidin and p53. Indeed, there is an increased nuclear:cytoplasmic ratio of p53 expression in binucleate *Cep55 −/−* NSCs and neurons compared to mononucleate cells **(Figure 9E)**. Furthermore, almost no mononucleate cells have p53 N:C ratios > 2, while more than 50% of binucleate progenitors and 20% of binucleate neurons do **(Figure 9F)**. These data suggest the existence of a p53-dependent pathway for apoptosis of binucleate cells in the cortex, that may be most sensitive in NSCs.

Our previous figures showed that apoptosis is increased in the brain but not body tissues of *Cep55* knockouts. *Cep55 −/−* MEFs have defective ESCRT recruitment during abscission, and increased binucleation. Therefore, we asked whether they also had increased p53 expression or apoptosis. Surprisingly, the answer appears to be no, neither apoptosis nor p53 levels are increased in *Cep55 −/−* MEF cultures **(Figure 9G-I)**. Furthermore, binucleate *Cep55 −/−* MEFs do not have any detectable difference in p53 expression compared to mononucleate MEFs **(Figure 9J)**. These data suggest the possibility that the p53-dependent apoptotic response to binucleation/tetraploidy, and/or its consequences, is regulated differently in various cell types, contributing to the dramatic tissue-level differences in phenotypic severity observed in germline *Cep55* knockouts.

### *Cep55* knockout forebrain apoptosis and size but not body size and longevity are p53-dependent

To test the hypothesis that p53 activation is the cause of apoptosis and microcephaly in *Cep55 −/−* mice, we crossed the *p53* knockout to the *Cep55* knockout. Mouse knockouts of *p53* have normal brain size and structure at birth, with almost no consequences for development (Insolera et al., 2014; Jacks et al., 1994). As expected, *Cep55 −/−* mice with wild-type p53 status exhibit microcephaly **(Figure 10A, second panel)**. We detected no difference in brain size with heterozygous deletion of p53 **(Figure 10A, third panel)**. However, complete deletion of p53 partially rescues brain size **(Figure 10A, right-most panel), and B, C)**. *Cep55;p53* double knockout cortical length is 21% longer than *Cep55 −/−*, but still 10% shorter than wild-type controls **(Figure 10B)**. Furthermore, cortical area is 32% larger than *Cep55 −/−*, but still 19% smaller than wild-type controls **(Figure 10C)**. Next, we evaluated whether deletion of p53 prevents apoptosis in the *Cep55* knockout. Indeed, apoptosis in *Cep55 −/−* mice is p53-dependent **(Figure 10D-E).** Therefore, preventing apoptosis partially rescues brain size in *Cep55 −/−* mice, but not to wild-type size. These results highlight the importance of Cep55 function for sufficient NSC proliferation, as even with apoptosis inhibition in *Cep55* knockouts brain growth is impaired.

**Figure 10.**
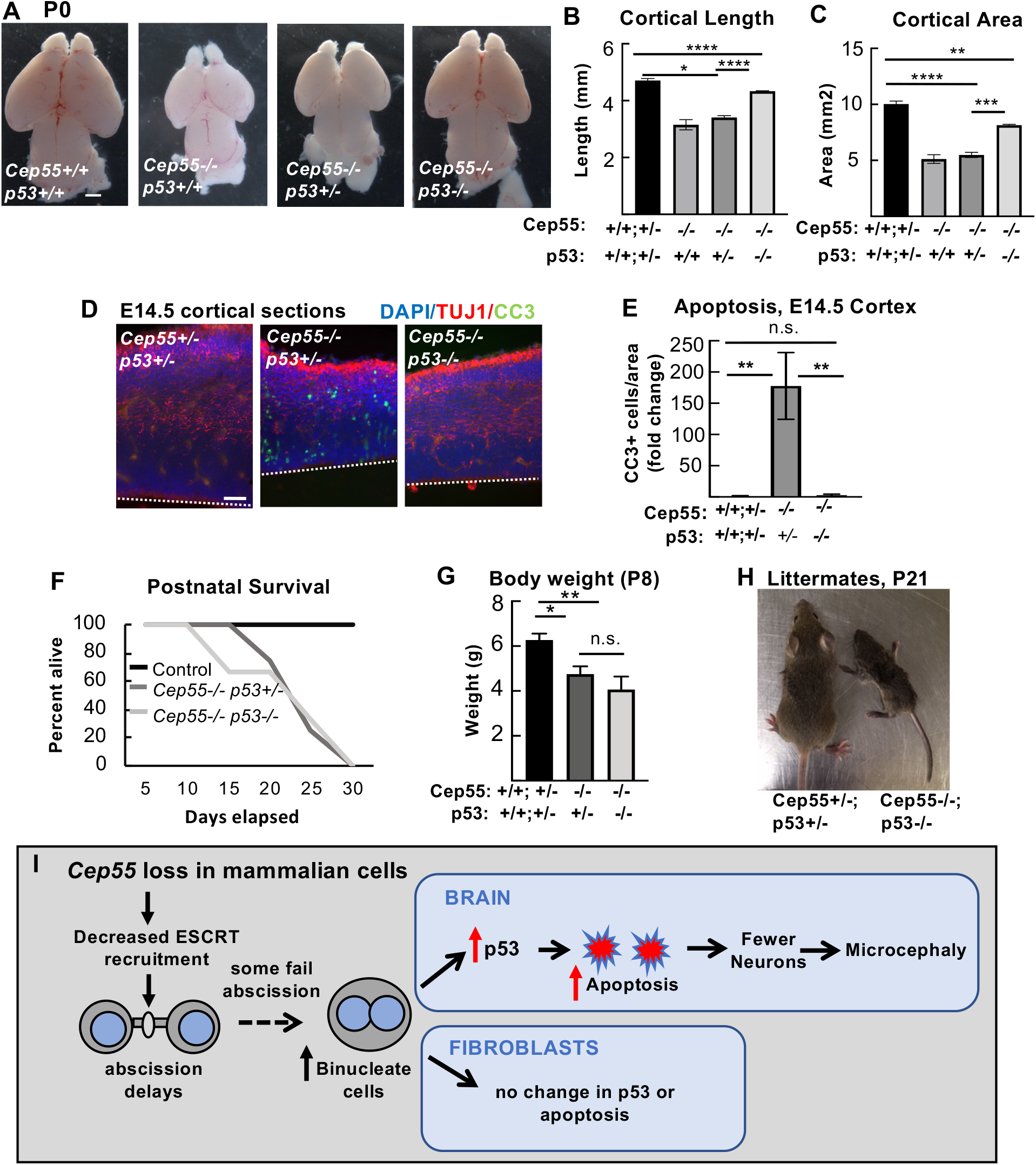
*p53* deletion blocks apoptosis and partially rescues brain size, but not body size or survival in *Cep55* knockout pups. **(A-C)** Co-deletion of *p53* partially rescues microcephaly in *Cep55 −/−* mice. **(B-C)** Homozygous *p53* knockout shows partial rescue of *Cep55−/−* cortical length and area. **(D-E)** Apoptosis of *Cep55−/−* cortex is fully blocked in *Cep55;p53* double knockouts. Dashed line in D = apical membrane. **(F-H)** *p53* co-deletion does not improve *Cep55−/−* pups' postnatal survival (F) or body weight (G,H). For B,C; n=11 controls (*Cep55+/+,+/−; p53+/+,+/−),* 2 *Cep55−/−;p53+/+*, 5 *Cep55−/−;p53+/−*, 3 *Cep55−/−;p53−/−.* For E, n=3 controls *Cep55+/+,+/−*; *p53+/+,+/−*, 2 *Cep55−/−;p53+/+*, 3 *Cep55−/−;p53−/−*. For F and G, n=15 controls, 4 *Cep55+/−;p53−/−* and 3 *Cep55−/−;p53−/−*. * p <0.05, ** p < 0.01, *** p < 0.001, **** p <0.0001, n.s., not significant. B,C,F: one-way ANOVA. E: one-way ANOVA, Tukey’s multiple comparisons test. **(I)** Working model for the effects of *Cep55* loss on abscission of mammalian cells, and for the differential effects on neural stem cells versus fibroblasts. *Cep55−/−* mouse cells have impaired ESCRT recruitment and abscission delay. Most cells complete abscission, but at least some fail to complete cytokinesis resulting in increased binucleate cells. Binucleate NSCs and neurons have elevated p53 expression, but not binucleate MEFs. p53 activation in NSCs and neurons causes apoptosis in Cep55 ko cortices, which partially accounts for microcephaly.

We earlier noted that *Cep55 −/−* mice suffer postnatal lethality starting in the second week of life, and have reduced body size at birth and poor postnatal growth **(Figure 1 – supplemental figure 1A, B)**. Therefore, we wanted to investigate whether lethality or body size are improved in *Cep55;p53 −/−* mice, as brain size is. Surprisingly, the postnatal survival curve of *Cep55;p53 −/−* mice is not improved compared to *Cep55 −/−* single knockouts **(Figure 10F)**. Furthermore, body weight and size are still severely reduced in *Cep55;p53 −/−* pups, similar to *Cep55 −/−* **(Figure 10G, H)**. Thus, it appears there are disparities in both the consequences of *Cep55* loss for development of different tissues in the animal, and in the degree to which p53 co-deletion rescues brain versus body size.

## Discussion

In this work we investigated the roles of the cytokinetic abscission regulator Cep55 in embryonic development, particularly in the brain. Data from cell lines had led to a general model in vertebrate cells that Cep55 was critical for the recruitment of ESCRT components to the midbody, to complete abscission. It was perhaps expected that knockout of Cep55 would cause early embryonic lethality, as knockout of another cytokinesis gene, mgcRacGAP does (Van de Putte et al., 2001). However, we find that while some cells fail abscission in the absence of Cep55, most succeed, and embryos develop and survive past birth. ESCRT recruitment is indeed severely reduced in most *Cep55 −/−* midbodies, but it is not abolished. We find Cep55 acts to ensure the speed and success rate of abscission **(Figure 10 I).** This function of Cep55 appears to be especially important in NSCs, because it is essential for proper brain growth. NSCs that fail abscission and become binucleate activate p53 and apoptosis, but embryonic fibroblasts do not. Consistent with this, p53 co-deletion to prevent apoptosis partially rescues *Cep55 −/−* brain size, but not body size. Our work emphasizes the need for understanding specialized requirements for abscission regulation in different types of stem cells and growing tissues, and tissue-specific apoptotic responses to abscission stress.

Knockout of *Cep55* enabled us to test the model that Cep55 is required to recruit ESCRT components to the midbody, via interactions with Alix and Tsg101 (Carlton et al., 2008; Carlton & Martin-Serrano, 2007; Lee et al., 2008; Morita et al., 2007; Stoten & Carlton, 2018) and that this recruitment is necessary for abscission completion. Abscission includes both membrane scission and microtubule disassembly to sever the intracellular bridge; ESCRT components, recruited by Cep55, are thought to be necessary to couple these processes together temporally, by directly mediating membrane scission and recruiting the microtubule-severing protein spastin (Guizetti et al., 2011; Steigemann et al., 2009). Our findings appear to challenge two aspects of this model. First, we find correctly localized ESCRT components in some *Cep55 −/−* midbodies, suggesting Cep55 is not the sole recruiter of ESCRTs to the midbody in vertebrate cells. Second, we find that without Cep55, microtubule disassembly during abscission does not fail but is only delayed. In HeLa cell cultures, Cep55 depletion caused greatly increased midbody index, and cells remained at midbody stage for hours before either regressing or finally succeeding (Fabbro et al., 2005; Zhao et al., 2006). By contrast, in NSCs *in vivo*, we only observed a small increase in midbody index, and only a slight delay in microtubule disassembly in *Cep55 −/−* brains. These findings suggest the intriguing possibility that microtubule disassembly and membrane scission are uncoupled in the *Cep55 −/−* mouse, and that membrane scission is more severely delayed. It is possible that microtubule severing proteins could be recruited through post-translation modifications of tubulin instead of via ESCRTs (McNally & Roll-Mecak, 2018). This could explain why we observe excess MBRs at the apical membrane in knockout brains: they could be tethered by a hollow membrane bridge. Alternatively, increased MBRs in *Cep55 −/−* mice could indicate a problem in MBR disposal or engulfment, as MBRs can bind to cell receptors and be internalized, influencing downstream cell polarity and fate (Peterman et al., 2019; Pollarolo et al., 2011; Singh & Pohl, 2014). Clearly, more work is needed to refine the general abscission model to determine the exact roles of Cep55 at the molecular level, the mechanisms that may recruit ESCRT proteins besides Cep55, and whether either membrane scission or microtubule disassembly can be achieved without ESCRT components.

Tissue-specific p53-dependent responses appear to be a major factor in the disproportionate consequence of Cep55 loss for brain size compared to the rest of the body. This is consistent with recent evidence for variable sensitivity of different murine developing tissues to p53 activation (Bowen et al., 2019), and differential p53-dependent responses to cell stress in specific cell contexts (Kastenhuber & Lowe, 2017). First, we find increased p53 expression and apoptosis in *Cep55 −/−* binucleate NSCs, but not binucleate MEFs. Second, apoptosis is increased in *Cep55 −/−* CNS tissues, but not in other embryonic body tissues. Finally, we show prevention of p53-dependent apoptosis partially rescues brain size in *Cep55 −/−* mice, but does not improve body size or survival. It is likely that in multiple *Cep55 −/−* cell types, abscission delays and failures hinder proliferation, resulting in decreased growth of body tissues, but that in NSCs they additionally result in p53-dependent apoptosis. The explanation for this might be that some tissues have a higher tolerance for abnormal cells. Indeed, some mammalian tissues, such as the liver and heart, have binucleate cells normally (Guidotti et al., 2003; Paradis et al., 2014); and syncytia development by incomplete cytokinesis is part of testis development (Greenbaum et al., 2011; Ren & Russell, 1991). The brain appears to have a low threshold for damaged cells to activate p53; several microcephalies involve p53-dependent apoptosis (Bianchi et al., 2017; Houlihan & Feng, 2014; Insolera et al., 2014; Little & Dwyer, 2019; Mao et al., 2016). Since cortical NSCs undergo many rounds of cell divisions to stochastically create lineage trees of different daughter cell types (Dwyer et al., 2016; Llorca et al., 2019), even low rates of errors early in development can have profound effects on neuron production and brain growth.

*Cep55* mutants serve as models for an emerging spectrum of human brain malformations. Human mutations in *Cep55* also cause severe defects in CNS development and surprising relative sparing of other tissues. Different alleles identified so far seem to cause a range of phenotypic severity. Null (nonsense) mutations result in lethal hydranencephaly, a fluid-filled skull with almost no brain tissue (Bondeson et al., 2017; Frosk et al., 2017; Rawlins et al., 2019). Compound heterozygotes for one nonsense allele and one missense allele have microcephaly, a small brain with normal structure. Finally, splicing mutations in *Cep55* result in micro-lissencephaly, a small brain with fewer sulci and gyri. The latter two classes of patients survived (Barrie et al., 2020). Unlike humans and mice with *Cep55* mutations, a zebrafish *Cep55* mutant has small eyes (Jeffery et al., 2015; Yanagi et al., 2019). The developing eye is a neuroepithelium with NSCs that are similar to those in the brain, so it is unclear why the human and mouse do not have small eyes upon *Cep55* loss. A possible explanation is that fish have larger eyes than forebrains, so their eye growth may require the abscission speed and accuracy that Cep55 ensures. Together these data suggest Cep55 may have evolved in vertebrates to help build bigger more complex nervous systems.

During the writing of this manuscript, another research group published the phenotype of a *Cep55* knockout mouse (Tedeschi et al., 2020). They similarly found *Cep55 −/−* mice have microcephaly disproportionate to body size reduction, an accumulation of binucleate NSCs, and apoptosis specific to neural tissues. However, we performed additional analyses that cause us to differ with some of their interpretations. First, we differ in that we find ESCRTs are sometimes recruited in *Cep55 −/−* cells. We are the first two groups to attempt to answer this question quantitatively, and more work is needed by cell biologists to determine if and how ESCRTs can be recruited without Cep55 in vertebrate cells. Second, we show specific abscission defects occur in *Cep55 −/−* cortices, not simply binucleate cells: we find abscission delay, midbody structural defects, and MBR accumulation in our *in vivo* brain analyses. Importantly, we are the first to show Cep55 is not required for microtubule disassembly during abscission. Future work will determine if *Cep55 −/−* NSCs are completing abscission without ESCRT components, or whether recruitment is simply reduced or delayed. Third, we do find evidence of a role for Cep55 in abscission of non-neural cell divisions, suggesting that Cep55 and the ESCRT system are not dispensable for non-neural cells. Finally, we identify contrasting p53 responses to cytokinetic defects in different cell types as a potential mechanism leading to disparate secondary consequences and tissue phenotypes. Our data suggest that cell division defects and apoptosis are related, with increased p53 expression occurring in binucleate *Cep55 −/−* cortical cells. Future work by many groups with complimentary approaches will be needed to understand Cep55 knockout phenotypes, why Cep55 evolved in vertebrates, and the complex regulation of cytokinesis in developing tissues.

## Materials and Methods

### Mice

Mouse colonies were maintained in accordance with NIH guidelines and policies approved by the IACUC. Embryos were harvested by cesarean section, and the morning of the vaginal plug was considered E0.5. Littermate embryos served as controls for all experiments and all parent and embryo genotypes were confirmed by PCR. The Cep55 allele (strain C57BL/6NCep55^em1(IMPC)Tcp^ was made as part of the KOMP2 phase 2 project at the Toronto Centre for PhenoGenomics for the Canadian Mouse Mutant Repository. It was maintained on C57BL/6 and 50/50% C57BL/6 and FVB/N background embryos, which were used for experiments. We verified the correct mutation in our mutants with DNA sequencing and RT-PCR (data not shown). p53 knockout (Trp53^tm1Tyj^) mice on C57BL/6 background were obtained from The Jackson Laboratory [(Jacks et al., 1994); JAX stock #002101 The Jackson Laboratory Bar Harbor, ME]. These mice were bred with 50/50% C57BL/6 and FVB/N Cep55 knockout embryos for creation of the Cep55;p53 mouse line. Cep55 heterozygous mice were crossed with both the mT/mG reporter line (JAX stock #007576, The Jackson Laboratory Bar Harbor, ME) and Sox2-Cre mice (JAX stock #0085454, The Jackson Laboratory Bar Harbor, ME(Hayashi et al., 2002; Muzumdar et al., 2007) to produce mice that express plasma membrane-localized GFP. Sex of embryonic mice were not noted as sex was not a relevant biological variable for these experiments. The specific ages of embryonic mice used is noted in figure legends for each experiment.

For survival analyses, pups were kept with their mother and littermates. Milk was noted in the bellies of Cep55 −/− and Cep55;p53 double knockout mice until at least P2. Pups were weaned at 21-25 days, unless too small to be separated from mothers. In some situations, pups were euthanized due to severe failure to thrive for humane reasons, and that was recorded as the day of death; in most cases pups died without intervention.

### Immunoblotting

MEF cells were washed with ice cold PBS, and lysed directly on the plate (6cm) into lysis buffer (500 μl); 50mM Tris HCl, 150mM NaCl, 1% NP-40, 1mM EDTA, 1mM DTT, 1mM PMSF, 1X HALT protease/phosphatase inhibitor cocktail. After incubating lysates on a rotisserie at 4° C for 30 minutes, lysate were cleared by centrifugation at 21k x g for 15 min at 4° C. Supernatants were diluted to 1x with 5X Laemmli buffer for western blot. Equivalent amounts of cells within each experiment were run on a 4-20% polyacrylamide gel. After transfer onto nitrocellulose with the BioRad Transblot Turbo, membrane was blocked with Licor TBS blocking buffer for 1 hour at room temp. Primary and secondary antibodies were diluted in 50% blocking buffer, 50% TBS, final 0.1% Tween-20. Primary antibodies were incubated overnight at 4° C, and after washing, followed by appropriate species specific near-infrared (680 or 800nm) secondary antibodies for 1 hour at room temperature. Blots were imaged with the dual color Licor Odyssey CLx imager. Densitometry of western blots was performed with Licor Image Studio software. Primary antibodies used: rabbit monoclonal anti human beta-actin (1:10,000, Clone13E5 4970 Cell Signaling, Danvers, MA) and mouse monoclonal anti-mouse Cep55 raised against amino acids 163-462 mapping at the C-terminus (1:1000, sc-377018 Santa Cruz, Dallas, Texas).

### Immunostaining and H&E Staining

To collect cryosections for immunohistochemistry in supplemental figure 2, and figures 3, 5, 9, 10 and 11, age E12.5, E14.5 and P0 brains were removed from heads and fixed for 4, 6 and 24 hours, respectively, in 4% PFA, followed by immersion in 30% sucrose in PBS overnight. Whole E10.5 embryos (used in figure 9) were fixed overnight. Next, whole brains or embryos were embedded in O.C.T. compound (Tissue-Tek 4583, Sakuraus Torrance, CA) and cryosections were cut at 20 μm thickness and collected on Superfrost Plus slides (Fisher Scientific, 12–550-15). Frozen sections were stored at −80 degrees. Prior to immunostaining, cryosections were warmed to room temperature, then if antigen retrieval was needed (Pax6, Tbr2, p53 antibodies), immersed in 10 mM citrate buffer at 95 degrees for 20 minutes. After cooling, sections were blocked in 2% NGS for 1 hour, followed by incubation with primary antibodies overnight at 4 ° C. The next day, after PBS washes, sections were incubated with AlexaFluor secondary antibodies at 1:200 and DAPI at 1:100 for 30 min followed by PBS washes and coverslipping with VectaShield fluorescent mounting medium.

For immunofluorescence (IF) on coverslips of dissociated cortical progenitors and MEFs (figures 2, 6, 7, and 10), a similar protocol was used but with primary antibodies applied for 3 h at room temperature. Antigen retrieval was not used in dissociated progenitors except in the case of Tbr2 immunostaining; coverslips were immersed in 0.07M NaOH pH 13 for 2 min before permeabilization. Coverslips were mounted on Superfrost Plus slides with Fluoro-Gel (Electron Microscopy Sciences, Hatfield, PA, 17985– 10).

Paraffin-embedded brains were sectioned and stained with hematoxylin and eosin (H & E) by the UVA Research Histology Core for Figures 1 and Supplemental Figure 2.

### Cortical cell cultures

Cells were dissociated from E14.5 cortices following a protocol adapted from Sally Temple’s laboratory (Qian et al., 1998). The Worthington Papain Dissociation Kit was used to dissociate cells (Worthington Biochemical Corporation, Lakewood, NJ, Cat # LK003150). Cells were cultured in DMEM with Na-Pyruvate, L-Glutamine, B-27, N2, N-acetyl-cysteine and basic fibroblast growth factor (Final Culture Media). After 24 or 48 hrs, cells were fixed by adding an equal volume of room temperature 8% PFA (paraformaldehyde) for 5 min to cell media, followed by removal of media and addition of −20° cold methanol for 5 min.

### Mouse embryonic fibroblast (MEF) cultures

Embryos from timed pregnant females were collected at E14.5. Embryos were decapitated, internal organs were removed from body cavities and remaining tissue was triturated with a 1 ml pipette and incubated in 1ml of 0.25% trypsin-EDTA for 25min at 37°C. The tissue was further triturated and tubes were spun at 1000 rpm for 5 min and the medium was aspirated off. The pellet was resuspended in DMEM/20% FBS with pen/strep added. The resulting cells from each embryo were plated in two 10 cm plates in DMEM/10% FBS. The next day, media was replaced. When approaching confluence, cells were trypsinized and plated on fibronectin-coated coverslips, grown overnight, and fixed with 4% paraformaldehyde (PFA) for 10 minutes or cold methanol for 10 minutes at 24 or 48 hrs.

### Cortical slabs for apical membrane view

Cortical slabs were prepared as previously described (Janisch & Dwyer, 2016). The meninges and skull were removed to expose the brain in E12.5 and E14.5 embryos, followed by fixation with 2% PFA for 20 min. Next, apical slabs were made by pinching off cortices, flipping so that the apical surface was upright, and trimming to flatten the slab. Slabs were fixed for another 2 min with 2% PFA followed by blocking with 5% normal goat serum (NGS) for 1 h. Primary antibodies were applied for 1 h at room temperature and then moved to 4° overnight. The next day, after 3 times, 10-minute PBS (phosphate-buffered saline) washes, secondary antibodies and DAPI were applied at a concentration of 1:200 for 30 minutes. After two more 10–minute PBS washes, slabs were coverslipped with VectaShield fluorescent mounting medium (Vector Laboratories Inc., Burlingame, CA H-1000) and imaged. z-stack depth was 8–20 μm and step size was 0.5 μm.

### Flow Cytometry

Cells from E15.5 brains were dissociated using the Papain Dissociation Kit (Worthington Biochemical Corporation). Single-cell suspensions were obtained by filtering through a 40 μm filter (BD Falcon Cell Strainer, blue nylon mesh, catalog #352340). For propidium iodide (PI) staining, cells were resuspended in 500 μl PBS and added to 4.5 ml ice-cold 70% ethanol for at least 2 hours. Samples were stored at 4°C. Fixed cells were rinsed in PBS and resuspended in 1 ml solution containing 100 μg/ml RNase A, 0.1% Triton X-100 and 50 μg/ml PI, and incubated at room temperature for 30 minutes in the dark. For Ki-67 and DAPI analysis, single-cell suspensions of E15.5 brains (*n*=3 pairs of *Cep55* mutants and littermate controls) were fixed in 1.5% PFA for 15 minutes on ice. Cells were then washed twice with FACS buffer (2% BSA, 1 mM EDTA, 0.01% sodium azide, PBS) and permeabilized in 0.1% Triton X-100 in PBS for 30 minutes on ice. Cells were washed twice in FACS buffer and 2×10^6^ cells were incubated in 100 μl FACS buffer with 1 μg/ml DAPI and 2 μl anti-Ki-67 antibody (monoclonal rat anti-Ki-67 Alexa Fluor 647, clone SolA15, eBioscience, Waltham, MA). Following three washes with FACS buffer, fluorescence was measured using a FACSCanto II flow cytometer (Becton Dickinson, Franklin Lakes, NJ). At least 20,000 events were collected per sample. Data were analyzed using FlowJo software (TreeStar).

### Abscission Live Imaging and Analysis

This method was previously described in (McNeely & Dwyer, 2020), based on first establishing SiR-tubulin for abscission duration studies in HeLa cells (Janisch et al., 2018). Briefly, E13.5 slabs were prepared as described above and placed apical surface down in a glass-bottom dish (MatTek, Ashland, MA, P35G-1.0–20-C) containing 50 nM SiR-Tubulin (Cytoskeleton, Denver, CO CY-SC002) in Final Culture Medium (described above). Each dish was placed in a humidifying chamber and into a 37 °C incubator with 5% CO2 overnight (approximately 15 h). The next day, the cortices were removed from the incubator, and prepared for imaging by adding Matrigel (Corning, Corning, NY, 356237; 1:3 dilution in Final Culture Medium) and a coverslip (Fisher 12–545-100) over the top of the cortical slab explant. Matrigel was allowed to solidify for 5 min in the incubator before final culture medium with 2% 4-(2-hydroxyl)-1-piperazineethanesulfonic acid (Gibco, Waltham, MA 15630080) was added to the dish, and then, imaging was performed. An Applied Precision (GE) DeltaVision with a heating plate and 40× objective (numerical aperture 1.53) was used for time-lapse image acquisition; z stacks of images ∼10-μm deep (z steps 0.4 μm) were taken every 15 min for abscission for up to 6 h. To minimize phototoxicity, the neutral density filter was set at 32% or lower, and the exposure time was kept to a minimum (<0.1 ms per slice). Deconvolved full z stacks and maximum intensity projection images were analyzed using ImageJ. For abscission duration, time 0 was midbody formation as ascertained by SiR-Tubulin appearance (compact microtubule bundles). Abscission completion was scored as the time point when there was complete removal of microtubules on a midbody flank ascertained when the SiR-Tubulin signal intensity decreased to background level. Midbody membrane scission was shown to be temporally coincident with midbody flank microtubule disassembly by several previous publications using differential interference contrast or phase imaging of cell lines in two-dimensional dissociated cultures (Elia et al., 2011; Lafaurie-Janvore et al., 2013; Steigemann et al., 2009); however here, we cannot rule out the possibility that the midbody plasma membrane might remain connected for some period of time after the microtubules are gone.

### Antibodies

Antibodies used in this analysis: mouse monoclonal anti-mouse citron kinase (1:100, CITK; 611367 BD Biosciences, San Jose, CA), mouse polyclonal anti-human Cep55 (1:00, H00055165-B01P Abnova, Taipei, Taiwan), mouse monoclonal anti-mouse Cep55 (1:100 for immunofluorescence experiments, sc-377018 Santa Cruz,, Dallas, Texas), rabbit polyclonal anti-human CC3 (1:250, 9661s Cell-Signaling, Danvers, MA), rat monoclonal anti-mouse Tbr2 (1:200, 14-4875, eBioscience (Thermo Fisher Scientific), Waltham, MA), rabbit polyclonal anti-mouse Pax6 (1:200, PRB-278P, BioLegend, San Diego, CA), mouse monoclonal anti-rat Aurora B kinase (1:300, 611082 BD Biosciences, San Jose, CA), rabbit monoclonal anti-human Aurora B kinase (1:100, ab2254, Abcam, Cambridge, MA), rat monoclonal alpha-tubulin (1:300, NB600-506, Novus Biologicals, Centennial, CO), rabbit monoclonal anti-human PH3 (1:200, 3458 Cell Signaling, Danvers, MA), chicken polyclonal anti-mouse Nestin (1:600, NES, Aves Labs, Davis, CA), rat monoclonal anti-human Ki67 (1:100, 14-5698 eBioscience, Waltham, MA), rabbit polyclonal anti-mouse p53 (1:500, NCL-L-p53-CM5p Leica Biosystems, Wetzlar, Germany), rat monoclonal anti-human Ctip2 (1:400, 18465 Abcam, Cambridge, MA), rabbit polyclonal anti-mouse Tbr1 (1:200, 31940 Abcam, Cambridge, MA), rabbit monoclonal Satb2 (1:200 Ab92446 Abcam, Cambridge, MA), rabbit polyclonal pericentrin (1:500, 92371, BioLegend, San Diego, CA), mouse monoclonal Tubb3 (Tuj1) (1:500, 801201, BioLegend, San Diego, CA), mouse monoclonal phospho-histone H3 (Ser10) (1:200, 9706, Cell Signaling, Danvers, MA), Phalloidin Oregon Green or 568 (1:50, 07466, Al2380 Invitrogen, Waltham, MA), chicken polyclonal anti-human alpha-tubulin (1:100, ab89984 Abcam, Cambridge, MA), mouse monoclonal Alix (1:100, SC-53538 Santa Cruz, Dallas, Texas), mouse monoclonal Tsg101 (1:100, SC-7964 Santa Cruz, Dallas, Texas), rabbit polyclonal CHMP2A (1:100, 10477-1-AP Proteintech, Chicago, IL), rat monoclonal Zo-1 (1:50, R26.4DC, DSHB, Iowa City, IA) and polyclonal rabbit anti Zo-1 (1:50 61-7300, rabbit, Invitrogen (Thermo Fisher Scientific), Waltham, MA). All antibodies were validated for the application used in multiple previous publications.

### Imaging and statistical analysis

Images used for data analysis in supplemental figures 2F-G, and figures 2C-E, 6, 9, 10 and 11 were collected on either a Zeiss Axio ImagerZ1 microscope with AxioCam MRm or a Zeiss AxioObserver fluorescent widefield inverted scope microscope. Images in figures 2A-B, 3, 4, 5A-B, 7 and 8 were taken on an inverted DeltaVision with TrueLight deconvolution microscope with softWoRx Suite 5.5 image acquisition software (Applied Precision (GE Healthcare), Issaquah, WA). A Leica MZ16F microscope with DFC300FX camera was used for images in Figures 1 and 11A-C, and supplemental figures 1 and 2A-E. Control and mutant fluorescence images were collected with the same exposure times and on the same day. All image analysis was performed in ImageJ/Fuji and any changes to brightness and contrast were applied uniformly across images. Statistical analyses were performed using Excel (Microsoft) or GraphPad PRISM software. The sample sizes were pre-determined based on our lab’s previous experience with cortical development analyses and others’ published results. After obtaining pilot data, power analyses were performed if necessary to determine if the number of samples obtained was high enough for the effect size seen. NSC cultures that were unhealthy were not imaged and analyzed, but no other data was excluded from the analysis. No randomization or blinding was used as no experimental manipulation was applied other than genetic knockouts. Genotyping was performed after collection of embryos to determine genetic status. Statistical tests used are specified in each figure legend. For each sample set a statistical test of normality was performed using GraphPad PRISM software. Parametric tests were used when sample sets had a normal distribution and non-parametric tests were used when sample sets did not have a normal distribution. Variance was calculated and was typically similar between groups.

## Acknowledgements

This work was supported by NIH Grants RO1 NS076640 and R21 NS1006162 to N.D.D., NIH Grant F30 HD093290 to J.N.L., and UVA Cell and Molecular Biology Training Grant 2T32GM008136-31A1 to J.N.L. We thank Bettina Winckler, Xiaowei Lu, Ann Sutherland, Michael McConnell, Todd Stukenberg and their laboratories for advice and discussion. We thank Adriana Ehlers, Gabrielle Wolfe and Naaz Daneshvar for help with cryosectioning and taking images. We are grateful to the Canadian Mouse Mutant Repository for use of the Cep55 allele sperm samples and to the UVA Genetically Enginereed Murine Model (GEMM) Core for in-vitro fertilization services. We thank the UVA Flow Cytometry Core for use of flow cytometry facilities.

---

**Figure 1 – figure supplement 1:**
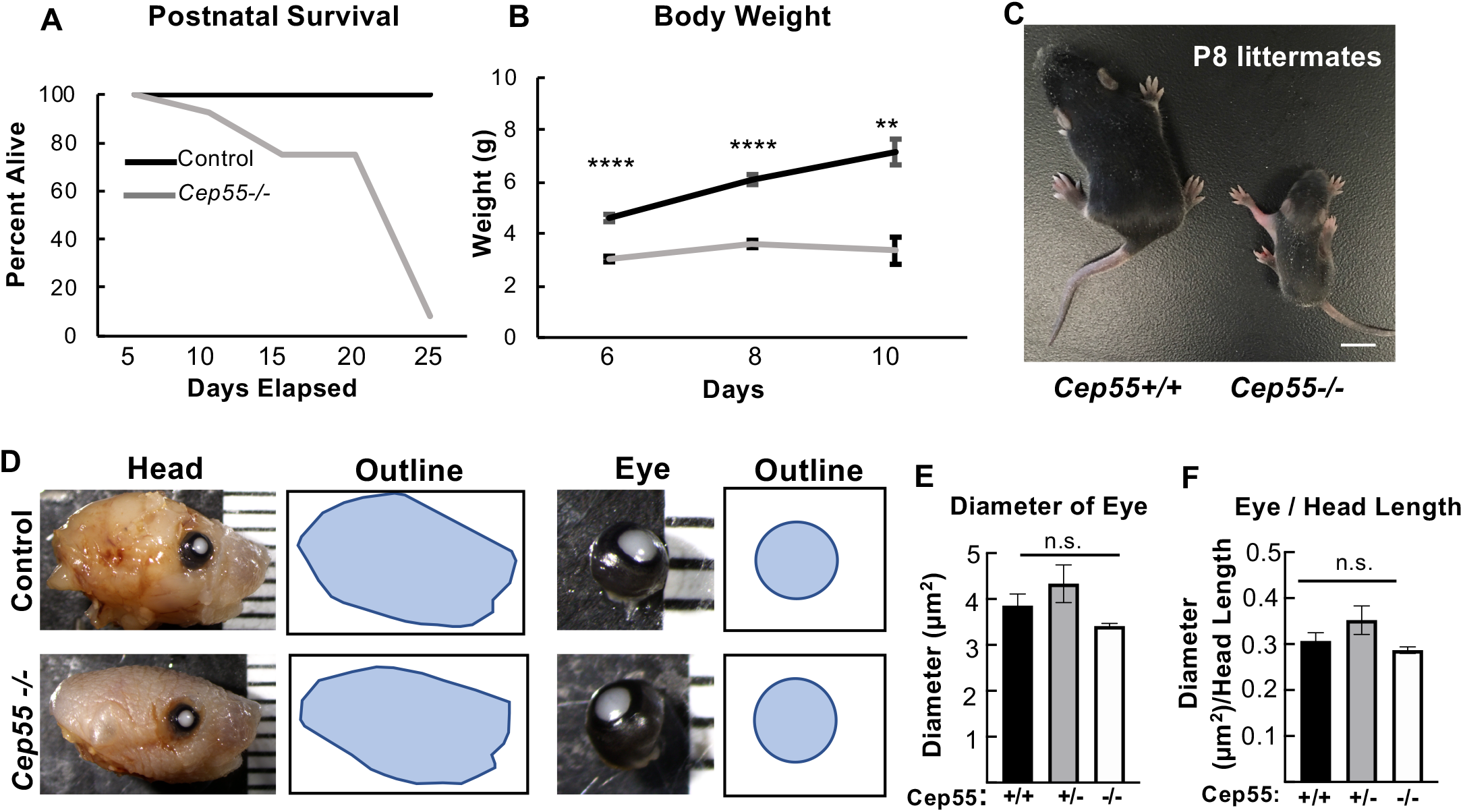
*Cep55* knockout mice exhibit normal eye size and can survive postnatally, but exhibit failure to thrive and peri-weaning lethality. **(A-C)** *Cep55 −/−* mice exhibit failure to thrive and pre-weaning lethality. (A) Most *−/−* pups die in the second or third week of life, and none survived past 25 days. n = 46 controls, 12 *Cep55−/−*. (B-C) *Cep55−/−* mice fail to gain weight between 6 and 10 days postnatally and are significantly smaller than controls. n, days: 6 = 13 controls and 6 *−/−*; 8 = 26 controls and 13 *−/−*, 10 = 11 controls and 4 *−/−*. Examples of postnatal day 1 control and *Cep55 −/−* heads and dissected eyes. Note flatter head shape of *Cep55 −/−* apparent in the outline. **(E)** Average eye diameter is not significantly changed in *Cep55 −/−* pups. (**F**) Normalizing eye diameter to length of head showed no change between control and *Cep55 −/−*. n= 6 +/+, 9 *+/−*, 5 *−/−* pups. Scale bar: C, 5 mm. n.s., not significant, ** p <0.01, **** p <0.0001. B: Student’s t-test, E, F: One-way ANOVA.

**Figure 1 – figure supplement 2:**
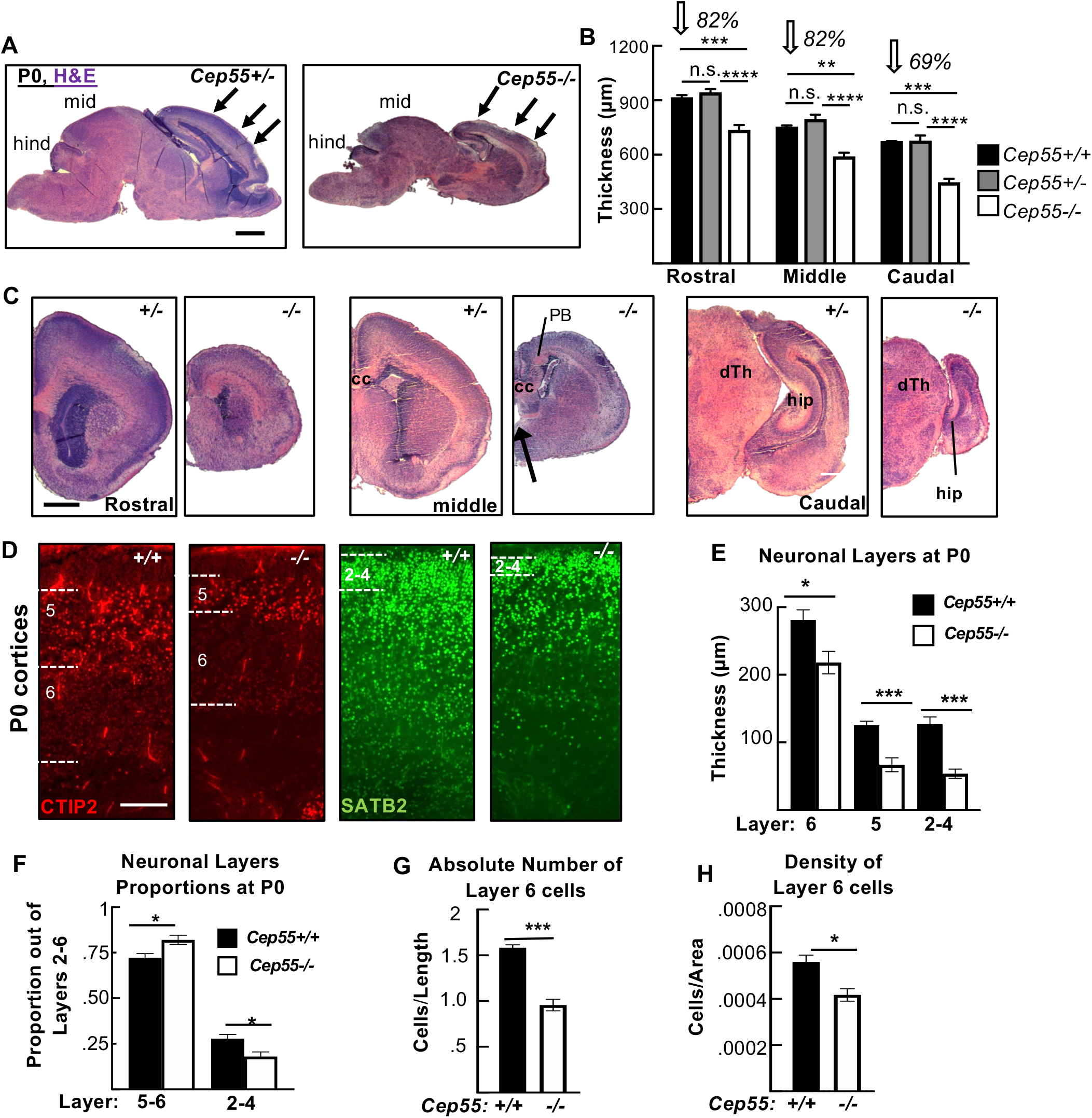
*Cep55* knockout cortices are thinner, especially in caudal cortex, with reduced thickness but preserved order of neuronal layers and reduced neuron density. **(A, C)** Representative *Cep55+/−* and *−/−* sagittal (A) and coronal (C) sections pf P0 brains stained with hematoxylin and eosin (H&E). Arrows in A mark approximate locations of rostral, middle,and caudal, coronal sections in C. mid and hind signify midbrain and hindbrain; PB, Probst bundles of axons; hip, hippocampus; cc, corpus callosum; dTh, dorsal Thalamus. **(B)** The mean cortical thickness in *Cep55−/−* brains is 82% of normal in rostral and middle sections, and only 69% of normal in caudal sections. For B, n = 3 Cep55+/+, 5 Cep55+/−, and 5 Cep55−/− mice. (**D)** Representative images of P0 *Cep55+/+* and *−/−* cortical sections labeled with antibodies to Ctip2, marking layers 6 (faint staining) and 5 (bright staining) and Satb2, marking layers 2-4. **(E)** All neuronal layers are significantly thinner in *Cep55 −/−* cortices at P0 **(F)** Layers 5-6 occupy a larger proportion of cp while layers 2-4 occupy a decreased proportion of the cp in *Cep55−/−* cortices compared to *+/+* controls. **(G)** The absolute number of layer 6 cells (# nuclei/length of cortex) is reduced by 40% in *Cep55−/−* cortices. **(H)** *Cep55−/−* layer 6 neurons are 25% less dense than wild-type cells (# nuclei/area). For G,H: n= 5 *Cep55+/+* and *4 Cep55−/−* mice; for I,K,J, n= 3 *Cep55+/+* and 3 *Cep55−/−* mice. Scale bar: A, C: 500μm, D: 100 μm. * p < 0.05; **p<0.01, ***, p<0.001, **** p < 0.0001. B: one-way ANOVA. G-J: t-test.

**Figure 3 – figure supplement 1:**
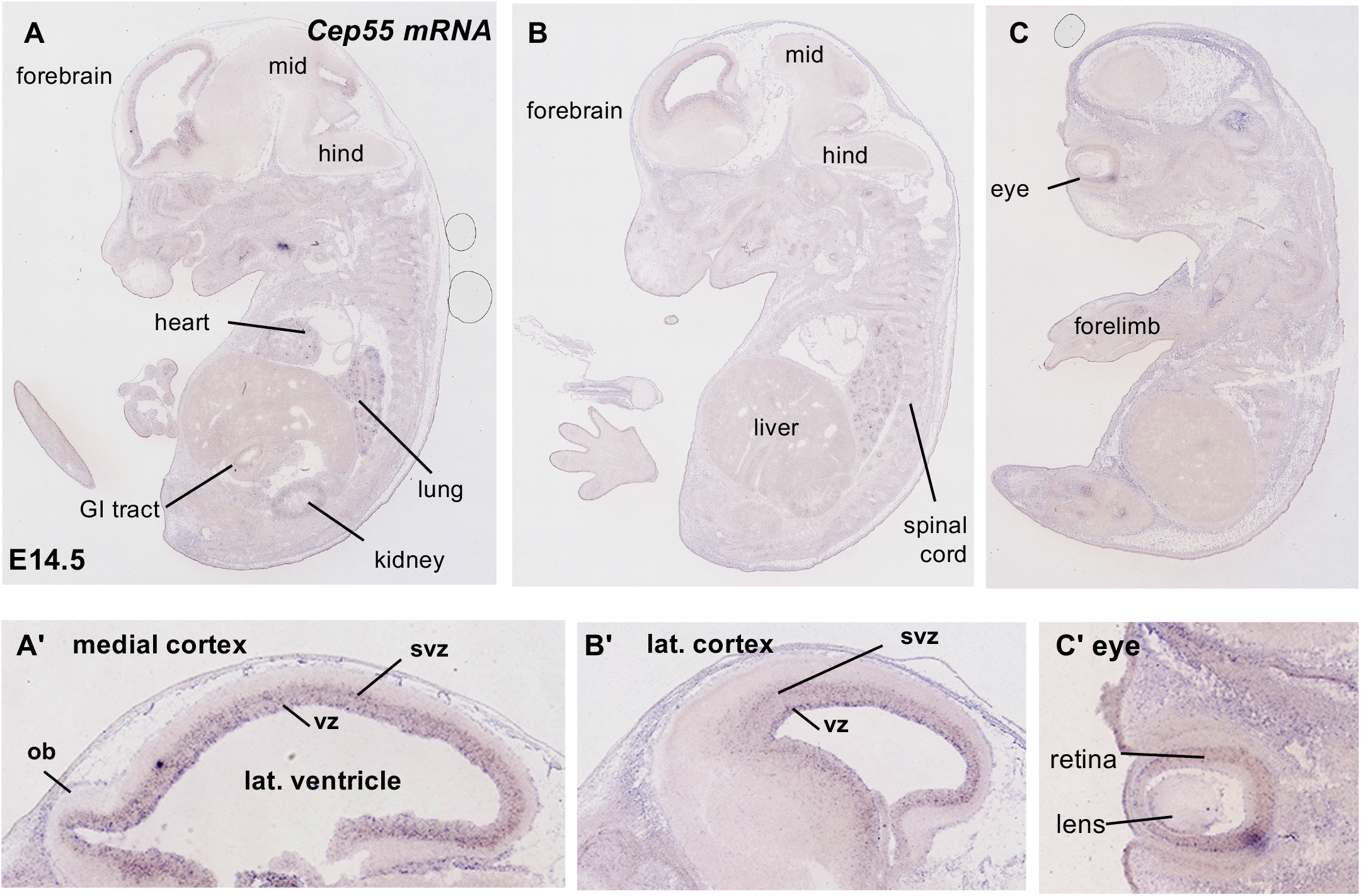
*Cep55* mRNA is expressed in proliferating embryonic tissues, including germinal zones of the nervous system. **(A-C)** Images of RNA in situ hybridization for *Cep55* mRNA on sagittal sections of E14.5 mouse embryos. *Cep55* is expressed in germinal zones of the nervous system including brain and eye, as well as widespread in proliferating embryonic tissues. Structures of interest in sagittal sections are labeled in whole embryo (A-C) and in zoomed forebrain (A’-B’) and eye (C’) images. mid: midbrain; hind: hindbrain; ob: olfactory bulb; vz: ventricular zone; svz: subventricular zone; lat vent: lateral ventricle. (Images from GenePaint, https://gp3.mpg.de, Visel et al. 2004).

